# Nucleotide-resolution Mapping of RNA N6-Methyladenosine (m6A) modifications and comprehensive analysis of global polyadenylation events in mRNA 3’ end processing in malaria pathogen *Plasmodium falciparum*

**DOI:** 10.1101/2025.01.07.631827

**Authors:** Cassandra Catacalos-Goad, Manohar Chakrabarti, Doaa Hassan Salem, Carli Camporeale, Sahiti Somalraju, Matthew Tegowski, Ruchi Singh, Robert W. Reid, Daniel A. Janies, Kate D. Meyer, Sarath Chandra Janga, Arthur G. Hunt, Kausik Chakrabarti

**Author notes:** Contributed Equally (Cassandra Catacalos-Goad, Manohar Chakrabarti).

## Abstract

*Plasmodium falciparum* is an obligate human parasite of the phylum Apicomplexa and is the causative agent of the most lethal form of human malaria. Although N6-methyladenosine modification is thought to be one of the major post-transcriptional regulatory mechanisms for stage-specific gene expression in apicomplexan parasites, the precise base position of m6A in mRNAs or noncoding RNAs in these parasites remains unknown. Here, we report global nucleotide-resolution mapping of m6A residues in *P. falciparum* using DART-seq technology, which quantitatively displayed a stage-specific, dynamic distribution pattern with enrichment near mRNA 3’ ends. In this process we identified 894, 788, and 1,762 m6A-modified genes in Ring, Trophozoite and Schizont stages respectively, with an average of 5-7 m6A sites per-transcript at the individual gene level. Notably, several genes involved in malaria pathophysiology, such as KAHRP, ETRAMPs, SERA and stress response genes, such as members of Heat Shock Protein (HSP) family are highly enriched in m6A and therefore could be regulated by this RNA modification. Since we observed preferential methylation at the 3’ ends of *P. falciparum* transcripts and because malaria polyadenylation specificity factor PfCPSF30 harbors an m6A reader ‘YTH’ domain, we reasoned that m6A might play an important role in 3’-end processing of malaria mRNAs. To investigate this, we used two complementary high-throughput RNA 3’-end mapping approaches, which provided an initial framework to explore potential roles of m6A in the regulation of alternative polyadenylation (APA) during malaria development in human hosts.

## INTRODUCTION

Post-transcriptional RNA modifications that make up an ‘epitranscriptome’, are emerging as important regulators of development and disease. Among diverse modifications, m6A is the most prevalent modification found within mRNAs that can affect multiple steps of transcript’s fate, including mRNA stability and translation (Meyer and Jaffrey 2014; Wang et al. 2014; Meyer et al. 2015; Wang et al. 2015). This methyl modification is a dynamic and reversible event driven by ‘writer’, ‘reader’ and ‘eraser’ proteins (Meyer and Jaffrey 2017). The coordinated action of methyltransferases (m6A writers: METTL3, METTL14 and other accessory proteins) and demethylases (m6A erasers: AlkB family members, FTO and ALKBH5) contributes to the deposition and depletion of this modification. In addition, YT521-B homology (YTH) domain family proteins act as “readers”(Liao et al. 2018) by directly binding to the N6-methyl group of m6A, thereby regulating diverse cellular functions (Meyer et al. 2015; Zhou et al. 2015; Mao et al. 2019; Meyer 2019b; Zhang et al. 2020). The biological consequences of m6A-mediated regulation are manifold, but a common theme is that they are required for many important aspects of mRNA processing and functions (Meyer et al. 2012; Meyer and Jaffrey 2014; Wang et al. 2014; Meyer et al. 2015; Wang et al. 2015; Bartosovic et al. 2017; Kasowitz et al. 2018). Particularly, m6A’s role in mRNA 3’ end processing was highlighted by several studies due to their preferential accumulation in the last exon, near stop codons or at the beginning of the 3’ UTR, which could mechanistically regulate polyadenylation/ alternative poly(A) site choice, RNA stability and translation (Dominissini et al. 2012; Meyer et al. 2012; Wang et al. 2014; Ke et al. 2015; Molinie et al. 2016; Bartosovic et al. 2017; Yue et al. 2018).

Messenger RNA 3’ ends, particularly 3’ untranslated regions (3’ UTRs), play crucial role in gene regulation by integrating multiple sequence-embedded signals to fine-tune gene expression. mRNA 3’ end formation – the process by which the primary transcript is cleaved and polyadenylated – is a near-universal feature of gene expression in eukaryotes. In yeast, animals, and plants, this process is mediated by a sizeable complex of some 15-20 protein subunits that recognizes a characteristic suite of sequence elements in the pre-mRNA (Tian et al. 2005; Chan et al. 2011; Di Giammartino et al. 2011). A number of broadly conserved cis elements regulate polyadenylation, the most central of which is an A-rich element (or family of motifs) situated some 15-30 nts 5’ (or upstream) of the actual site of processing and polyadenylation. In animals, this element is typified by the prevalent poly(A) signal AAUAAA (Proudfoot 2011); in yeast and plants, AAUAAA can function in this capacity, but other A-rich motifs have a similar functionality. Most eukaryotic pre-mRNAs have more than one potential PAS, and the choice of site often can vary during development or in response to environmental stimuli (Tian et al. 2007; Lutz and Moreira 2011; Xing and Li 2011; Elkon et al. 2013; Tian and Manley 2016). These variations can affect mRNA stability and function and help to tune overall transcriptional outputs and mRNA levels.

Evolutionarily, alternative 3’-end processing has been adapted to fit the needs of different organisms, contributing to the diversity of gene regulation and expression (Mayr 2016). Recent findings indicate intimate links between m6A methylation and mRNA 3’ end processing in deep branching parasitic protists, indicating early origin of m6A mediated regulation of mRNA metabolism in eukaryotic evolution. Specifically, apicomplexan parasites, such as *Toxoplasma gondii* among others uniquely contain a ‘YTH’ family m6A-reader as part of their machinery for 3’ end processing, cleavage and polyadenylation (Chakrabarti and Hunt 2015; Stevens et al. 2018; Farhat et al. 2021; Hunt et al. 2021). In *T. gondii*, functional WTAP, METTL3, and METTL14 subunits of the m6A writer complex proteins were identified and m6A peaks were detected in close proximity to the 3’-end of transcripts (Holmes et al. 2021). Further, loss of m6A resulted in aberrant transcriptional products in *T. gondii* from faulty transcription termination (Farhat et al. 2021), implicating a co-transcriptional role of m6A in mRNA processing. In the Kinetoplastid parasite *Trypanosoma brucei*, m6A methyltransferase writer complexes have not been identified (Balacco and Soller 2019). However, 50% of m6A marks were identified in the poly(A) tail of the actively expressed essential genes encoding for variant surface glycoproteins (VSGs), which appeared to stabilize the transcripts(Liu et al. 2019; Viegas et al. 2022). While these findings greatly underscore the importance of m6A biology, there is a lack of high-resolution data on transcriptome-wide RNA m6A methylation in parasitic protists. Also, it remains to be fully understood how the methylation pattern in transcripts provides protective responses for parasites in host environments via modulating post-transcriptional and translational control that are advantageous to parasites.

*Plasmodium falciparum* is an apicomplexan parasite which causes a lethal disease - malaria. In *P. falciparum*, m6A deposition is developmentally regulated during the human asexual stages of the parasite’s life cycle (Baumgarten et al. 2019) (Catacalos et al. 2022). In this parasite, m6A is deposited by the homologs of METTL3 and METTL14 methyltransferase proteins in *P. falciparum*, known as PfMT-A70 and PfMTA-70.2 respectively, and by WTAP proteins(Baumgarten et al. 2019; Govindaraju et al. 2020; Catacalos et al. 2022). However, no ‘eraser’ proteins have yet been identified. There are two m6A ‘reader’ proteins in *P. falciparum*. One of these is expressed exclusively in the Intraerythrocytic developmental cycles (IDC) (PfYTH.1: PF3D7_1419900)(Baumgarten et al. 2019; Govindaraju et al. 2020; Sinha et al. 2021) is the apicomplexan ortholog of the polyadenylation complex subunit CPSF4 (or CPSF30 (Stevens et al. 2018)); the presence of a YTH domain in a core polyadenylation complex subunit is also seen in plants but not mammals or yeast. The observation that a core subunit of the polyadenylation complex possesses an m6A -reader domain, lends itself to the hypothesis that m6A marks may be an important determinant of the 3’ end processing in *P. falciparum*. Interestingly, unlike the parasitic protists Trypanosoma and Toxoplasma sp., the *Plasmodium falciparum* genome is extremely A+T rich (∼90%)(Gardner et al. 2002). Therefore, it is possible that the deposition of N6-methyladenosine (m6A) epitranscriptomic mark may provide a higher-order signaling for processing mRNA 3’ ends in the exceedingly A-rich sequences in this organism, potentially triggering differential 3’-end processing via alternative polyadenylation (APA) (Yue et al. 2018; Hou et al. 2021; Chen et al. 2022). However, the nature of the polyadenylation sites and the contributions that APA make to gene expression, in *Plasmodium* species have not been extensively studied.

To resolve the above knowledge gaps, we provide here a nucleotide-resolution map of m6A epitranscriptome in *P. falciparum.* We used the DART-seq (Meyer 2019a; Tegowski et al. 2022) approach to accurately map m6A sites at single-nucleotide resolution in three different asexual developmental stages of *P. falciparum* in human RBCs, namely Ring, Trophozoite and Schizont stages, and successfully validated these m6A signatures using m6A-immunoprecipitation and high-throughput RNA sequencing, MeRIP-seq (Meyer et al. 2012). This base-resolution information should help to functionally characterize m6A’s role in parasite biology. Further, to test our hypothesis that m6A is a necessary component of the *P. falciparum* 3’ end processing machinery, we used two complementary genome-wide 3’ end mapping techniques to characterize poly(A) sites and APA in *P. falciparum*, which revealed that cleavage and polyadenylation occurs predominantly within sequence tracts that are exceedingly high in Adenosine (A)- content. Then, we compared the distribution of m6A at or near the 3’ end of mRNA coding regions and in the context of poly(A) sites. Together, these studies clarify many issues regarding transcriptome-wide m6A modification stoichiometry and its association with *P. falciparum* polyadenylation and they raise new and interesting questions regarding the role of RNA modifications in the processing of mRNA 3’ ends in this clinically important human parasite.

## RESULTS

### High-resolution mapping of N6-methyladenosines (m6As) in *P. falciparum*

Initial characterization of m6A modifications in the *P. falciparum* coding regions over the course of intraerythrocytic development showed that m6A is developmentally regulated in *P. falciparum* (Baumgarten et al. 2019; Govindaraju et al. 2020; Sinha et al. 2021) and the methylation occurs primarily within the eukaryotic consensus motif for m6A modifications, DRACH, particularly in the context of an enriched RAC (R= G/A) sequence (Baumgarten et al. 2019). However, the role of m6A modifications in malaria biology remains obscure, particularly because the existing m6A mapping data in *P. falciparum* localizes m6A residues within a range of 100–200 nt-long regions of transcripts instead of a precise base location.

To determine precise m6A modification sites in *P. falciparum* mRNAs, we used a nucleotide resolution m6A mapping technique, DART-seq, which identifies m6A modifications primarily in the context of the <u>RAC</u> motif (Meyer 2019a; Tegowski et al. 2022). DART-seq utilizes a fusion protein consisting of the m^6^A-binding YTH domain fused to the C-to-U deaminase APOBEC1, which targets cytidines adjacent to m^6^A sites for C-to-U editing. Using this approach, we mapped m6A in *P. falciparum* mRNA across three tightly synchronized intraerythrocytic developmental stages – Ring, Trophozoite and Schizont. Our analysis revealed variation in the methylation levels of several *P. falciparum* genes across the three asexual stages. In this process, we identified 8,983, 6,077, and 10,895 m6A sites in the Ring, Trophozoite, and Schizont stages, respectively, which corresponded to 894, 788, and 1,762 m6A-modified genes in each of the above stages. Among these sites, a significant portion are unique m6A modification sites, occurring in only one of the above three asexual developmental stages at any given time (**Fig. 1A**). These sites were further filtered to show enrichment strictly in RAC (R=G/A) motifs (**Fig. 1B**). An over-representation analysis, shown as a heatmap in **Fig. 1C**, revealed statistically significant Gene Ontology (GO) terms associated with genes that are involved in RNA metabolic processes, peptide biosynthetic pathways and oxidation-reduction processes. Dynamicity of m6A modifications were further analyzed on a nucleotide-to-nucleotide basis for each individual gene from this DART-seq analysis. A detailed IGV browser view of two representative mRNAs, Pf3D7_0418200, eukaryotic translation initiation factor 3 homolog, and Pf3D7_0418300, a conserved protein of unknown function, illustrate widespread occurrence of m6A in the coding region of these transcripts (**Fig. 1D**). Importantly, we observed significant overlap between all m6A sites identified by DART-seq (**Fig. 1D**, ‘all’) and those that occurred strictly within the RAC motif (Fig. 1D, ‘RAC’) for several genes in all three RBC developmental stages (Ring, Trophozoite and Schizont). Some of these DART-seq defined sites are also within the coverage peaks previously identified by Baumgarten et. Al. (2019) (Baumgarten et al. 2019) (**Fig 1D**, ‘m6A Peaks’, blue bars). As seen in these two genes, the majority of m6A sites are located near the 3’ end of the coding region, a trend which is common in many *P. falciparum* mRNAs. Further, DART-seq revealed that the extent of m6A modification in only one specific intraerythrocytic stage decreased from Schizont to Trophozoite/Ring stages. This is due to the occurrence of an extensive set of m6A sites across the mRNA sequence reads in the Schizont stage compared to the number of m6A containing genes with unique m6A locations found in the complete RNA sequence reads of the Ring and Trophozoite stages (**Supplemental_File S6**).

**Figure 1.**
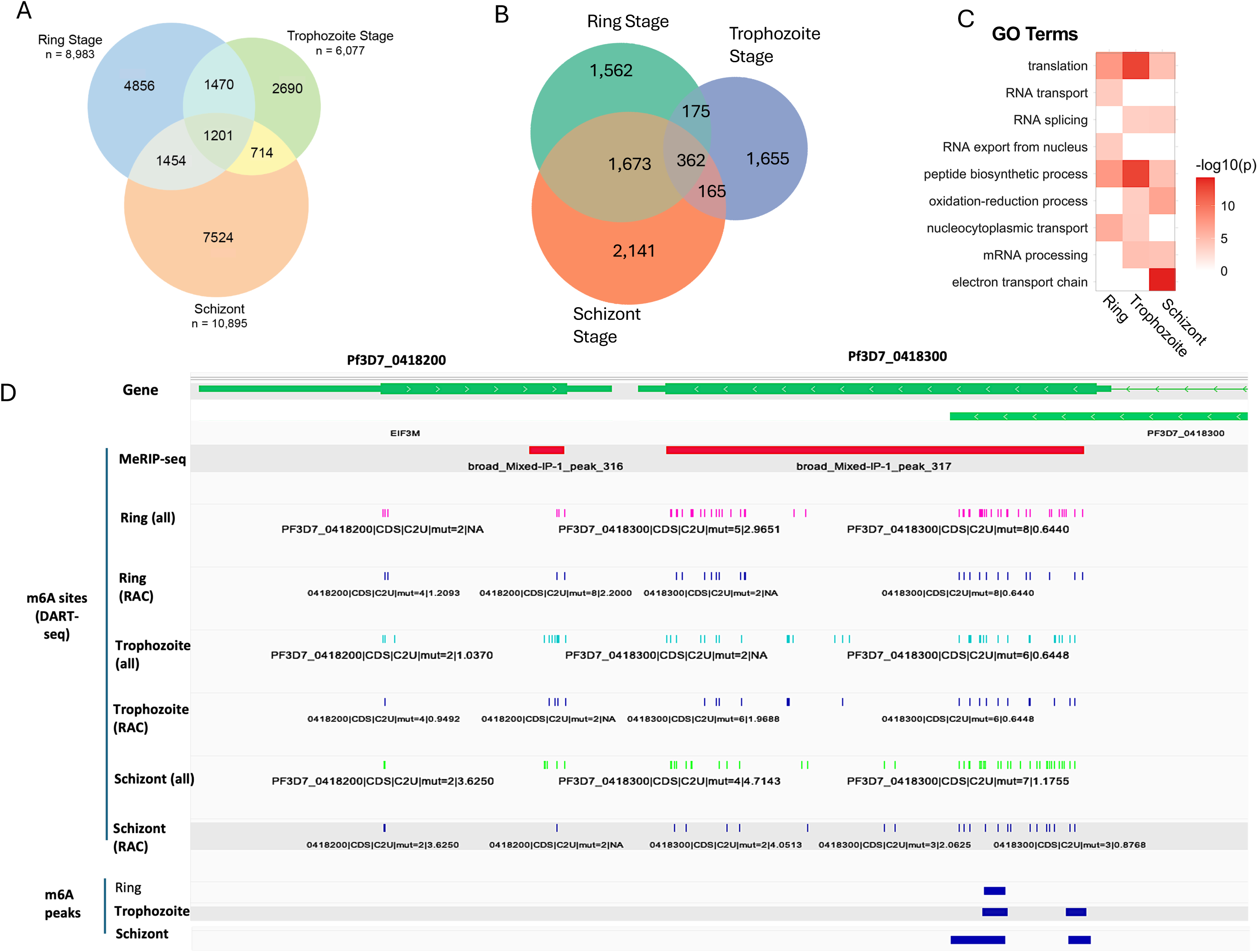
(A) Venn diagram showing the number of unique m6A modification sites between different *Plasmodium falciparum* asexual blood stages, i.e. Ring, Trophozoite and Schizont stages. (B) Venn diagram showing the number of unique m6A modification sites that are in the context of ‘RAC’ motif. (C) Significantly enriched GO terms across the three stages among genes containing m6A sites. (D) IGV screenshot showing data overlaps from nucleotide-resolution DART-seq (vertical lines) and meRIP-seq (red bar) from this study, and one that was previously reported (green bars) (Baumgarten et al. 2019) for a representative *P. falciparum* gene PF3D7_1035500 (MSP6).

Moreover, we compared the number of m6A modification sites per gene between different *P. falciparum* asexual blood stages (**Fig. 2A**), which showed most concordance between Ring and Trophozoite stages (R= 0.85, p<0.0001) whereas correlation of methylation patterns in Trophozoite and Schizont stages turned out to be more stochastic (R= 0.69, p<0.0001). Genes with a difference in m6A site counts greater than 30 between stages are highlighted as outliers in blue, suggesting potential stage-specific regulation for these genes. Next, we determined the distribution of m6A modification sites across normalized gene length for the above three plasmodium life stages, which were categorized by their locations within the gene: whole gene, 5’ end (0-0.33 interval of gene length), middle (0.33-0.66), and 3’ end (0.66-1.0). A Kruskal-Wallis test was conducted for each category to assess the differences in m6A distribution across the three life stages (**Supplemental_Fig_S1**). Analysis of the general distribution of m6A patterns in the gene body revealed a prevalence near the stop codon or near the 3’ end/ last exon of the transcript. The quantitative landscape of m6A in *P. falciparum* for unique modification sites is also represented by box plots in **Fig. 2B** across normalized gene length with respective P-values (**Supplemental_File_S5**). We observed a strong preference for m6A methylation in the 0.66-1.0 intervals of the gene length corresponded to the 3’ end of the genes approaching stop codons (about 3-4-fold increase compared to other regions of transcripts). Genes which are uniquely enriched in m6A in a stage-specific manner in the *P. falciparum* transcriptome were also analyzed (**Supplemental_File_S6**). The list of m6A unique genes from each *P. falciparum* stage were plugged into Cytsoscape ClueGo application (G. Bindea and H. Fridman) to obtain the enriched ontologies and pathways at high confidence (p<0.05). Enrichment observations from this analysis are visualized in **Supplemental_Fig_S2**, **S3** and **S4** for Ring, Schizont and Trophozoite developmental stages, respectively. As a result, we annotated a large set of m6A-containing genes with known GO terms that were enriched in biological processes of RNA transport, proteosomal function, mRNA processing, oxidation-reduction processes, and others. Additionally, the frequency of m6A unique genes along with their products across the three plasmodium developmental stages are depicted in a heatmap in **Supplemental_Fig_S5**.

**Figure 2.**
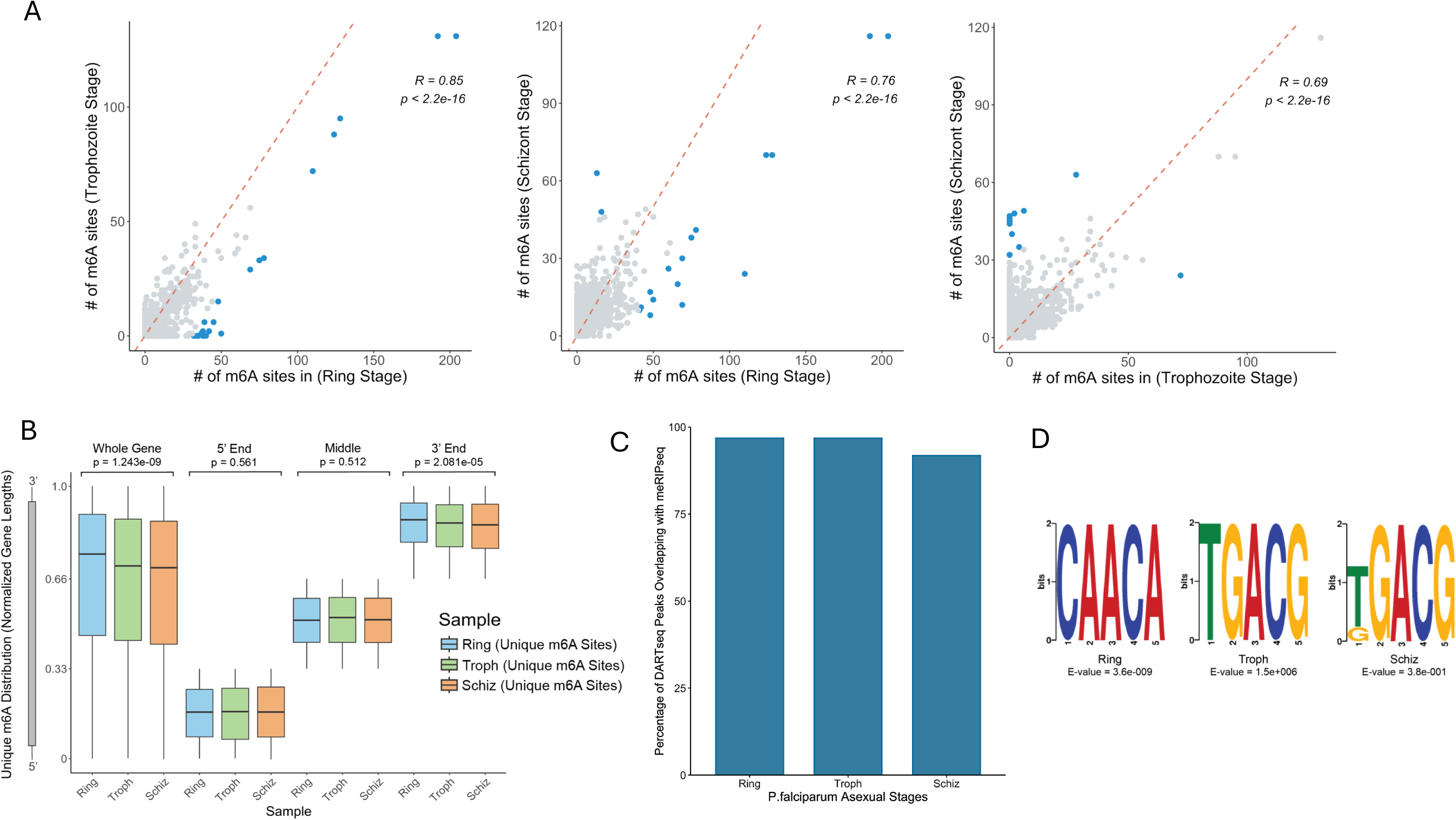
(A) Scatterplots showing the number of m6A modification sites between different Plasmodium falciparum life stages. Plots compare (left to right) Ring vs Trophozoite, Ring vs Schizont, and Trophozoite vs Schizont stages. Data points representing genes with a difference of more than 30 m6A sites between the two compared stages are highlighted in blue. Spearman correlation coefficient and p-value are marked on the figure. (B) Box plots show the distribution of unique m6A modification sites across normalized gene length for above three *P. falciparum* asexual RBC stages in the human host. (C) Percentage of DART-seq sites which are also detected from mixed-stage meRIP-seq sites. (D) Stage-specific variation and frequency of DRACH motifs utilized in three different asexual blood stages of *P. falciparum*. Motif discovery was conducted using MEME (Bailey et al. 2015)

To validate the m6A sites identified by DART-seq, we conducted an overlap analysis with MeRIP-Seq data, which offers a broader genomic perspective on m6A site distribution in *P. falciparum*. MeRIP-Seq was performed with immunoprecipitated RNA obtained from asynchronous, mix stage *P. falciparum* using an anti-m6A antibody. We hypothesized that the m6A transcripts that are identified by DART-seq from three intraerythrocytic developmental stages of *P. falciparum* should be proportionately represented in these m6A-immunoprecipitated samples. We observed a robust 92-97% overlap among the three *P. falciparum* life-stages (**Fig. 2C**), affirming the precision of DART-seq in detecting m6A modifications at a single nucleotide resolution.

To further substantiate the m6A sites identified by DART-seq, publicly available *P. falciparum* m6A data (Baumgarten et al. 2019) was incorporated. Overlapping these peaks with our in-house MeRIP-Seq data revealed a 55%-58% overlap, which could be due to the much lower number of samples in the publicly available m6A data. This is consistent with the finding that DART-seq identified numerous methylation sites per mRNA, as shown in **Fig. 1D**. While m6A MeRIP-seq data (**Fig. 1D**, red horizontal bar) intersects almost entirely with DART-seq peaks for these transcripts, m6A peaks from a previous study (Baumgarten et al. 2019) corresponded only partially to DART-seq peak locations (**Fig. 1D**, blue horizontal bars, ‘m6A Peaks’). This underscored the fact that DART-seq profiling facilitated robust m6A quantification at nucleotide resolution in *P. falciparum* asexual stages. **Supplemental_Files_S5** and **S6** lists all m6A sites, including unique sites, identified by DART-seq in three *P. falciparum* RBC stages. Interestingly, several variations of the consensus DRACH motif were identified in the Ring, Trophozoite and Schizont stages (**Fig. 2D**). The E-value is calculated based on its log likelihood ratio, which reflects how significantly different the observed motif is compared to a random sequence, with higher log likelihood ratios indicating a stronger match to the motif model. The three motif logos across the stages are representative of the significantly enriched patterns containing the RAC consensus sequences in mRNAs surrounding the m6A peaks.

### m6A enriched RNAs are involved in stress response and various metabolic processes important for parasite biology

The malaria parasite *P. falciparum* completes its asexual development following invasion of human RBCs, in about 48 hours. During the asexual stage development, gene expression in malaria parasite is tightly regulated and the rate of transcription and decay varies throughout the parasite’s life cycle (Llinas et al. 2008; Hollin and Le Roch 2020). Throughout the blood-stage infection, these parasites are constantly exposed to a range of potentially damaging extracellular stimuli and therefore, should elicit survival responses. However, the regulatory mechanisms for these responses are largely unknown. For example, invasion of merozoites in uninfected RBCs mark the beginning of early asexual developmental stage (ring stage), which requires parasites to survive in febrile temperatures - a fundamental biological property critical for the parasite’s successful propagation in the human host and persistent infection(Tinto-Font and Cortes 2022). Recent transcriptomic studies have revealed that the protective heat-shock response in *P. falciparum* requires modulation of expression of numerous heat-shock responsive genes and is intimately linked to artemisinin resistance (Tinto-Font et al. 2021; Zhu et al. 2022). From our DART-seq data, we have identified several of these stress response genes that are enriched in m6A sites. For example, the transcripts of canonical heat shock family proteins, such as the cytosolic PfHSP70 (PF3D7_0818900 and PF3D7_0708800) and PfHSP70x (PF3D7_0831700) (Grover et al. 2013), all possess extensive m6A methylation. Additionally, the major cytosolic *P. falciparum* HSP90 (PF3D7_0708400), which is known to play an essential role in the development of the parasite, particularly forming a chaperone complex with PfHSP70 to coordinate the folding of proteins important for the development of the parasite (Dutta et al. 2022), is also heavily m6A methylated in all three RBC stages of this parasite (**Fig. 3A**).

**Figure 3.**
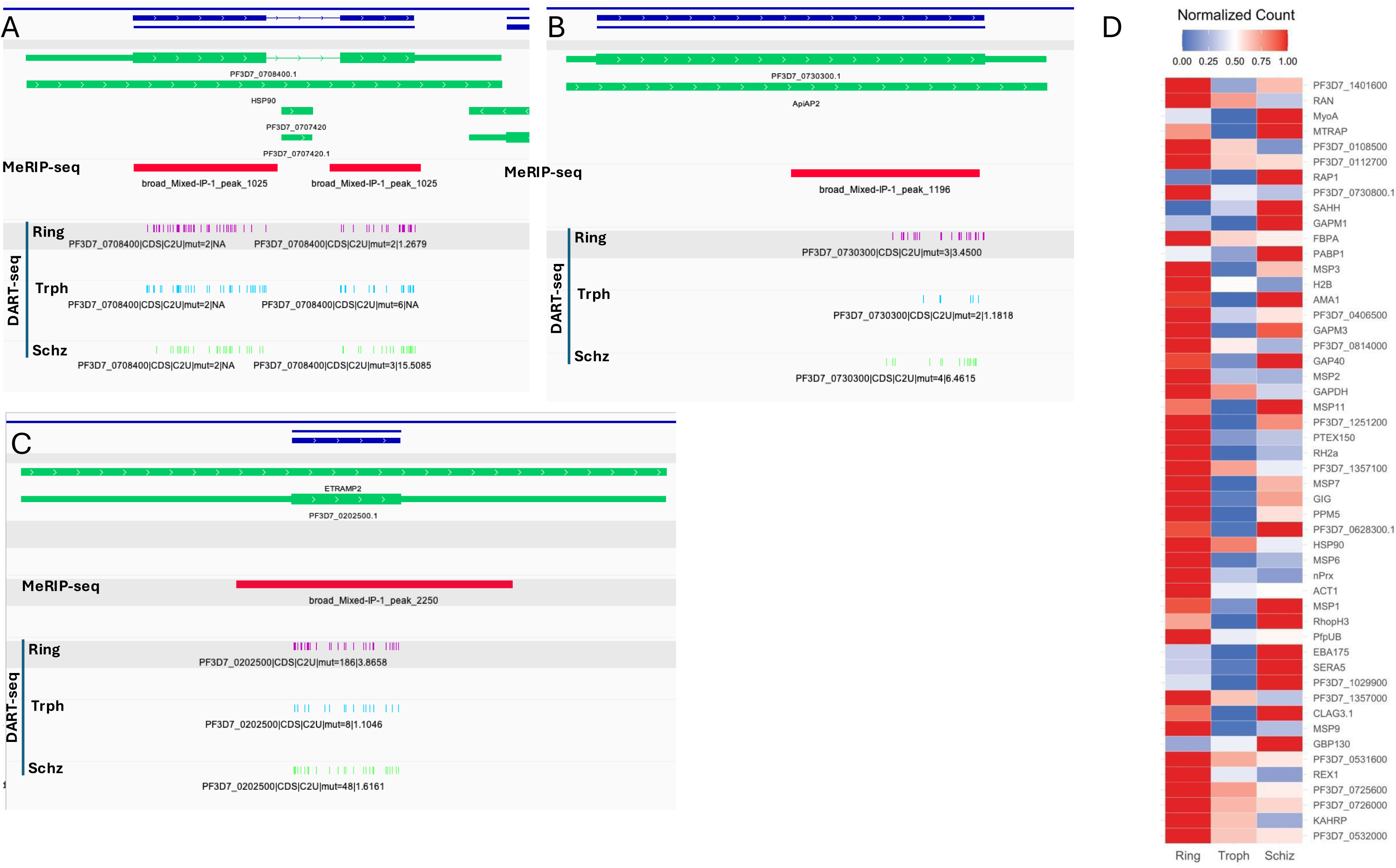
Dynamic landscape of m6A modification of mRNAs and Noncoding RNA: (A) Heat Shock Protein 90 (HSP90), (B) ApiAP2-L and (C) ETRAMP2. (D) Top 50 most m6A enriched genes and their differential methylation pattern in *P. falciparum* Ring, Trophozoite and Schizont stages.

Transcriptional gene regulation in *P. falciparum* is driven to a great extent by the Apicomplexan APETALA2 (ApiAP2) family of DNA-binding proteins(Campbell et al. 2010) that are homologous to plant transcription factors and have no mammalian counterparts. Our DART-seq and MeRIP-seq methods identified selective members of the ApiAP2 transcription factor family in *P. falciparum* that contain significant m6A marks in their transcripts. Particularly, m6A methylation is abundantly present in ApiAP2 member Pf3D7_0420300 mostly in the Ring stage whereas significant m6A enrichment was observed in all three asexual RBC stages in AP-2L(Pf3D7_0730300) (**Fig. 3B**), the ApiAP2 family member essential for the liver stage development of *P. falciparum*. In addition, some of the ApiAP2 proteins contain modest levels of m6A, such as in Pf3D7_0802100, Pf3D7_0622900, Pf3D7_1143100 and Pf3D7_0934400, as evident from the DART-seq data.

In addition, *P. falciparum* post-transcriptional and translational apparatus appeared to be substantially m6A modified. For example, several RNA helicases, such as ATP-dependent RNA helicase - DDX6 (DOZI)(Mair et al. 2006) that plays a role in the sexual development and translational repression in *P. falciparum*, UAP56 (Pf3D7_0209800) and DDX47 (Pf3D7_0218400), ribosomal proteins- S11 (Pf3D7_0317600), S26 (Pf3D7_0217800), L13 (Pf3D7_0214200), P2 (Pf3D7_0309600), S19 (Pf3D7_0422400), L31 (Pf3D7_0503800) and others, eukaryotic translation initiation factor 4E homolog (Pf3D7_0315100), translation enhancing factor (Pf3D7_0202400), elongation factor 1-delta (Pf3D7_0319600) and small nuclear ribonucleoprotein family, SNRPD and SNRPF and all the core RNA components of ribosome machinery, both small and large subunit rRNAs, are m6A modified. Interestingly, none of the components of the m6A epitranscriptomic machinery, such as the m6A writer methyltransferases or reader proteins are methylated, indicating a lack of epitranscriptomic autoregulation of the m6A machinery.

The membrane structure of a malaria parasite is important for many aspects of the parasite’s pathophysiology. Two unique membranous features that play essential roles in malaria biology are the parasitophorous vacuole membrane (PVM)(Goldberg and Zimmerberg 2020) and Maurer’s cleft (MC)(Tilley et al. 2008; Maier et al. 2009). The interface between parasite and host cell plays a key role in propagation and pathogenicity of intracellular *P. falciparum*, which is provided by the PVM, whereas MCs are novel membranous structures established by the parasite that concentrate virulence protein factors for delivery to the host cells. MCs are thought to originate through budding from the PVM or from extensions of the PVM termed the tubovesicular network. Two PVM family proteins, EXP1 and ETRAMPs, are essential for nutrient uptake across the PVM as well as maintaining architectural framework for the PVM(Spielmann et al. 2006; Mesen-Ramirez et al. 2019). EXP1 gene (PF3D7_1121600) shows abundant m6A modifications in all three coding exons. Among the ETRAMPs, ETRAMP2 (Pf3d7_0202500) transcripts are extensively m6A modified (**Fig. 3C**). Together with the above two proteins, m6A marks in the subtilisin-like protease 1(Collins et al. 2013) (SUB1, Pf3D7_0507500; methyl marks identified only in Ring and Schizont stages), which activates rapturing the parasite’s PVM and host cell membrane to continue erythrocyte invasion and other life stages, suggest that parasite’s protein, nutrient and metabolite export(Desai et al. 1993) and egress mechanisms are potentially regulated at the epitranscriptomc level. Additionally, the Serine Repeat Antigen (SERA) gene family, which is expressed in the asexual blood stages in *P. falciparum* and localized in the parasitophorous vacuole is abundantly modified with m6A marks.

*P. falciparum* critically relies on precise regulation of several antigens during asexual blood stage developments. For example, the *P. falciparum* knob-associated histidine-rich protein (KAHRP)(Maier et al. 2009) forms membrane protrusions that allows the major *P. falciparum* protein antigen PfEMP1 to concentrate and mediate cytoadherence, which in turn helps the parasite to escape host immune responses(Waller et al. 1999). Thus, KAHRP renders a pivotal role in malaria cytoadherence, the major virulence mechanism. Among all the *P. falciparum* genes that are m6A modified, KAHRP (Pf3D7_0202000) transcripts are most heavily methylated in all three asexual stages (**Supplemental_Fig_S6A**), suggesting epitranscriptomic control in the gene expression. In addition, transcripts for almost all the members of the Merozoite Surface Proteins (MSPs)(Beeson et al. 2016), which are predominantly composed of resident glycosylphosphatidylinositol (GPI) anchored proteins (MSP1-7, 9, 11 and DBLMSP) contain moderate to high m6A marks. Because these proteins are displayed abundantly over the merozoite surface, the free-living blood stages of malaria that can mediate the primary attachment stages for parasite invasion after rapture of schizont stage, several of these proteins, such as MSP1, MSP3 and MSP4 are vaccine targets for malaria. KAHRP, MSP1 and other genes, which make the list of top 50 most m6A enriched genes and their differential methylation pattern in *P. falciparum* Ring, Trophozoite and Schizont stages, is shown in the heatmap in Fig. 3D.

### Spatial organization of Poly(A) sites in malaria transcriptome determined by PAT-seq and TE-seq

The genomic distribution of cleavage and polyadenylation (poly(A)) sites can be reliably quantified at nucleotide resolution using RNA-seq. Earlier, Siegel et al. (Siegel et al. 2014b) described a collection of poly(A) sites in *P. falciparum* that were identified by mining RNA-Seq data, extracting poly(A)-containing reads, and mapping these to the *P. falciparum* genome. To understand the nature and content of poly(A) sites from this existing dataset and to compare it with Poly(A) site data from other Apicomplexan parasites (Farhat et al. 2021; Hunt et al. 2021), the nucleotide compositions surrounding the 3443 major poly(A) sites described in *P. falciparum* dataset (Siegel et al. 2014b; Siegel et al. 2014a) (provided in **Supplemental_File_S7**) were analyzed. For this, the genomic coordinates for each site reported were extended in both directions by 100 nts, and the position-by-position nucleotide compositions computed and displayed. The results (**Fig. 4A**) showed a decided enrichment for Adenosine (A) between 15 and 20 nts upstream from the poly(A) site, an enrichment above and beyond the remarkably high A content of sequences surrounding these sites. The region from 10 nts upstream to 10 nts downstream was from the actual cleavage site was enriched in U, except for a distinct peak of A adjacent to the poly(A) site. Throughout these regions, G and C content was very low, characteristic of the *P. falciparum* genome in general.

**Figure 4.**
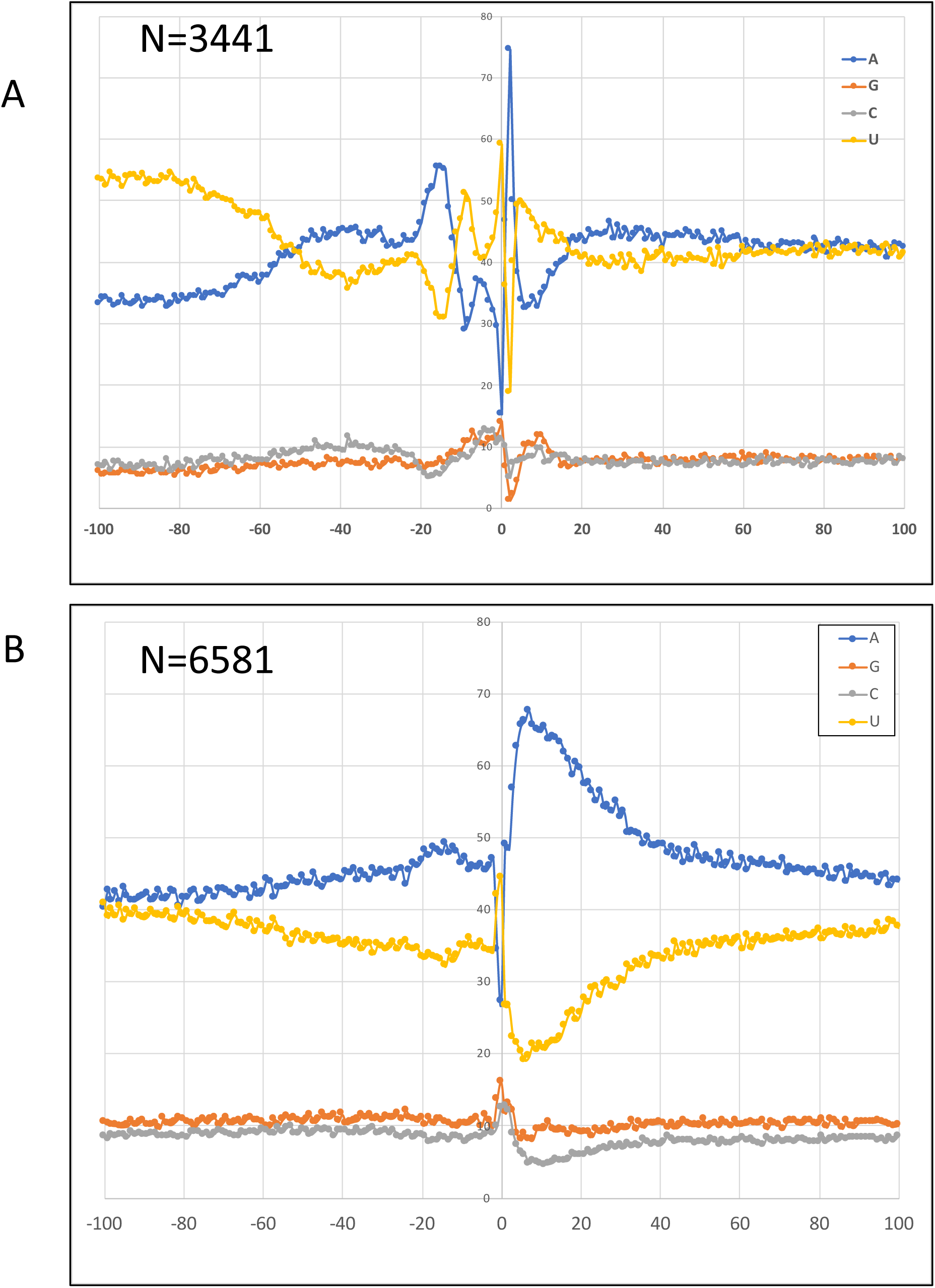
Nucleotide composition surrounding *P. falciparum* poly(A) sites. Sequences extending 100 nts upstream and downstream from the poly(A) sites listed in - (A) Siegel et. al. (Siegel et al. 2014a) or (B) Yang et al. (Yang et al. 2021) were collected and analyzed for overall base content (on a position-by-position basis).

More recently, long-read sequencing of full-length cDNAs was used to generate a collection of *Plasmodium* polyadenylation sites by Yang et. al. (Yang et al. 2021). These sites were collected into a dataset (**Supplemental_File_S7**) and the nucleotide compositions surrounding these sites determined. The results (**Fig. 4B**) were distinctive from the profiles shown in **Fig. 4A**. In particular, besides a high A content through the region surrounding the poly(A) site, there is little additional structure to be seen. In addition, in contrast to the content map shown in **Fig. 4A**, the apparent poly(A) sites in **Fig. 4B** are flanked on their 3’ ends by regions of very high A content. This corroborates with the predominance of A nucleotides at or surrounding poly (A) sites in other Apicomplexan single-celled parasite of cattle, *Babesia bovis (Li and Du 2014)*.

Given the differences seen in **Fig. 4A** and **4B**, two additional and complementary approaches to study poly(A) sites were applied to the task of poly(A) site characterization in *P. falciparum*. These approaches focus on the production of libraries that query just the RNA 3’ ends (as opposed to full-length RNA-seq as was done in the studies cited in the preceding paragraphs).

One approach was the so-called PAT-seq approach described elsewhere (Wu et al. 2014; Bell et al. 2016). For this, RNA was isolated from *P. falciparum* asexual growth stages in human RBCs, namely Ring (R), Trophozoite (T) and Schizont (S) stages and short cDNA tags that query the mRNA-poly(A) junction (Poly(A) Tags, or PATs) were generated and sequenced. Importantly, the reverse transcription (RT) reactions used anchored oligo-dT primers, much as was done for the long-read sequencing (Yang et al. 2021). This approach thus should corroborate the results described in Yang et al. (Yang et al. 2021). The sequences were trimmed to remove the residual oligo-dT tract present in the reverse transcriptase primer and mapped to the *P. falciparum* genome. These results were used to generate a list of poly(A) sites (**Supplemental_File_S8**).

Similar to the analysis performed for the poly(A) sites in Figs. 4A and 4B, the regions between 100 nts upstream to 100 nts downstream from each site defined by PATs were analyzed. The results (**Fig. 5A**) were similar to those obtained by long-read RNA sequencing (Yang et al. 2021). Specifically, there was a decided trend toward higher A content in the region within 50 nts upstream of these sites as well as a striking elevated A residue content immediately adjacent and downstream from these sites (**Fig. 5A**). This suggests that cleavage and polyadenylation may occur within tracts of A residues in the pre-mRNA. However, this result may also be explained by a large preponderance of internal priming by reverse transcriptase in the course of library preparation; even though the primer used in these experiments possessed a terminal VN dinucleotide, and should thus anchor the oligo-dT tract to the poly(A)-mRNA junction, it is conceivable that some amount of internal priming could occur, such that many of the sites assigned here may be internal stretches of A and not authentic 3’ ends.

**Figure 5.**
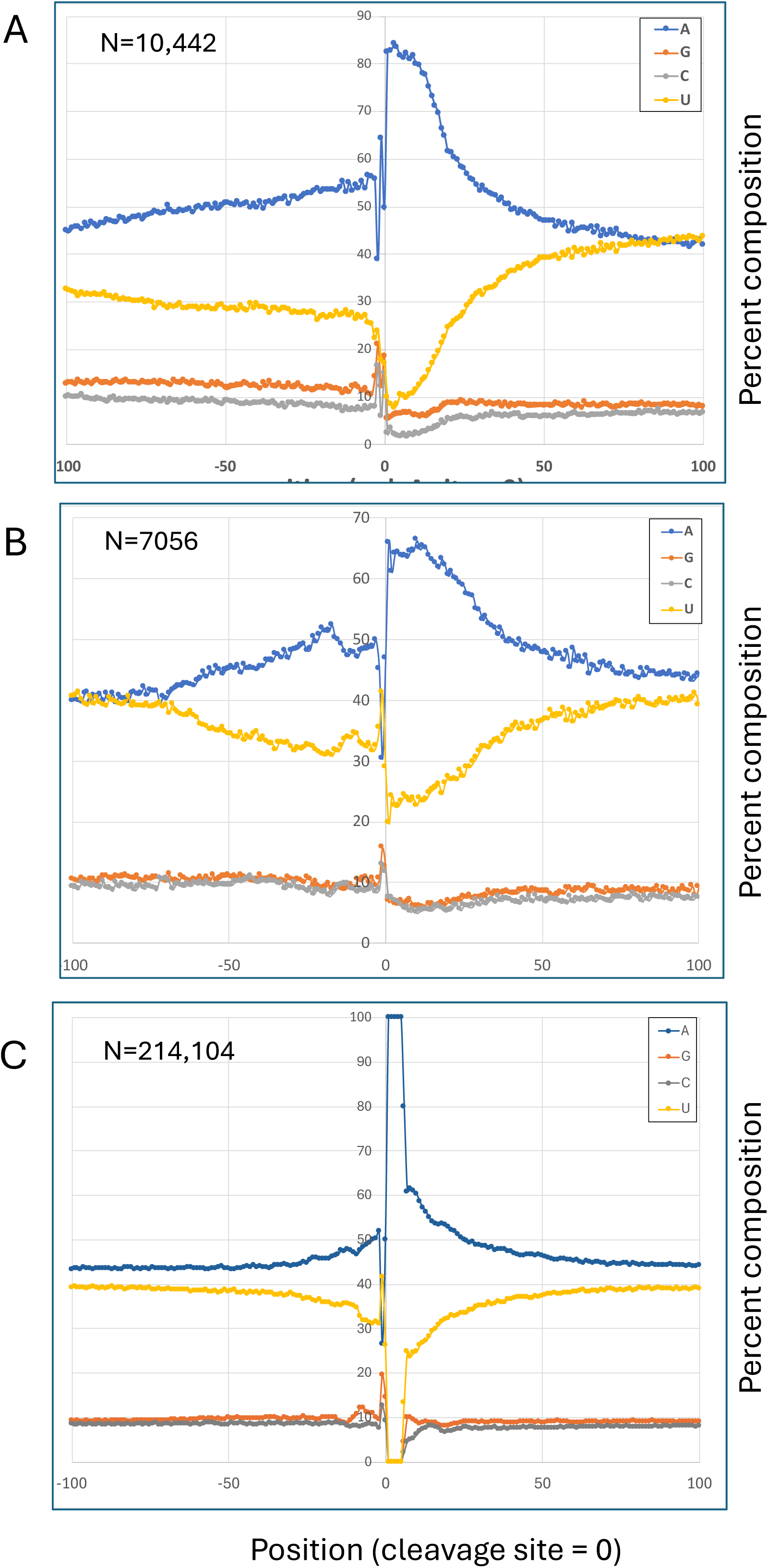
Nucleotide distribution surrounding poly(A) sites determined using (A) PAT-seq and (B) TE-seq. (C) Nucleotide compositions surrounding A-rich tracts in the *P. falciparum* genome.

To address this issue of internal priming, 3’-end libraries produced without oligo-dT priming were generated and sequenced (the process is illustrated in **Supplemental_Fig_S7**) using a unique approach, referred here as TE-seq. In order to do this, it proved necessary to identify and remove ribosomal RNAs as well as truncated (and often polyadenylated) rRNA fragments; this was necessary because a large number of the polyadenylated reads generated for the study shown in **Fig. 5A** mapped to rRNAs (**Supplemental_File_S2**). These RNAs, as well as unadenylated stable RNAs, would likely dominate the overall sequencing output for any such library, limiting the information that could be expected from such an effort. To remove these RNAs, an RNAse H-mediated approach was taken, wherein total RNA was treated (after ligation of a dedicated RNA adaptor) with RNAse H in the presence of a mixture of DNA oligonucleotides that were complementary to the 3’ ends of the expected interfering RNA species. In this way, the bulk of the unwanted RNAs could be separated from the RNA adaptor, which serves as the site for priming by reverse transcriptase.

For TE-seq, samples prepared using oligo-dT-independent priming (termed hereafter as TE-seq libraries) were sequenced and the reads processed and mapped to *the P. falciparum* genome. For this, reads that had at least 4 T’s adjacent to the RNA adaptor were extracted; this was done to recover reads that represent the poly(A)-mRNA junction. Subsequently, a list of 3’ ends that are defined by these reads was generated (**Supplemental_File_S9**). Using the TE-seq reads, analyses analogous to those shown in **Fig. 4** and **5A** were conducted. Akin to what was seen with the sites defined by PAT-seq reads, sites defined by TE-seq reads seemed to fall within extended regions of elevated A (elevated over and above the general high A content of these regions), with a pronounced peak immediately downstream from the apparent poly(A) site (**Fig. 5B**). In addition, there were hints of a region of very high A content between 15 and 20 nts upstream from the poly(A) site.

The studies presented in **Fig. 4B** and **Fig. 5** (**A** and **B**) collectively indicate that 3’ end processing of mRNAs in *P. falciparum* may occur within A-rich regions in the pre-mRNA, and that poly(A) tracts in many cases may include genome-encoded as well as posttranscriptionally-added bases. To further elaborate on this and rule out unanticipated artifacts arising from the AT-rich nature of the *P. falciparum* genome, the nucleotide compositions surrounding A-rich tracts in the genome were analyzed (**Fig. 5C**). As expected (given the high AT content of the *P. falciparum* genome), the position immediately adjacent to the 5’ end of the tract of A’s was more likely to be a U than either G or C (**Fig. 5C**). 3’ of the beginning of the tract, the A content was very high, and returned slowly (over the span of 40-60 nts) to the genome-wide averages for A residue content. These features are different from those seen in profiles surrounding polyadenylation sites defined by TE-seq.

### Stage-specific alternative polyadenylation in *P. falciparum*

As is seen in other systems, the collections of poly(A) sites assembled using either TE-seq or PAT-seq were such that most *P. falciparum* genes possess more than one site (**Fig. 6A and 6B**). This raises the possibility of alternative poly(A) site choice, such as in the different stages of growth and development of the organism. To explore this, differentially-utilized poly(A) sites were identified using the PAT-seq dataset, much as described previously (Bell et al. 2016; de Lorenzo et al. 2017; Chakrabarti et al. 2020). In this analysis, three pairwise comparisons were conducted between *P. falciparum* asexual RBC developmental stages: Ring vs. Schizont, Ring vs. Trophozoite, and Schizont vs. Trophozoite. For the Ring vs. Schizont and Ring vs Trophozoite comparisons, 562 and 537 individual poly(A) sites were differentially-utilized (**Supplemental_File_S10**); these affected 330 and 325 genes, respectively (**Supplemental_File_S10**). More than half of the genes in each set were shared in the other (**Supplemental_File_S10**), indicative of commonalities in the developmental impact on APA. Using the same stringent cut-off for the Schizont vs Trophozoite comparison, no sites were differentially-utilized (**Supplemental_File_S10**); with a more permissive cut-off, 54 sites affecting 38 genes were differentially-utilized.

**Figure 6.**
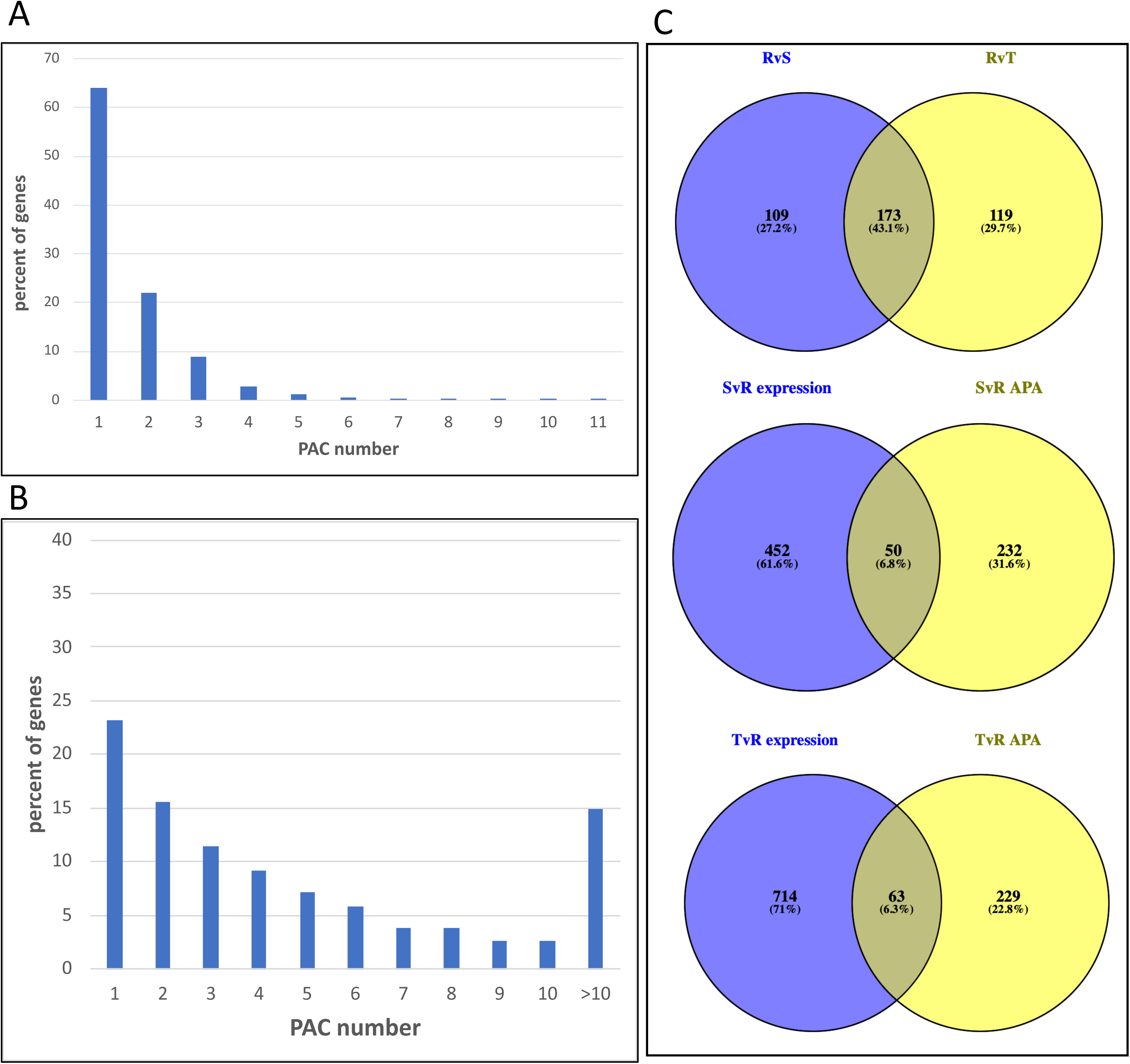
Measurement of poly(A) clusters per gene (PAC/gene) identified by (A) TE-seq (B) PAT-seq. These results illustrate the numbers of genes with multiple poly(A) sites, and thus can be affected by APA. (C) Venn diagrams of Genes affected by APA, particularly overlapping differential gene expression in Ring, Trophozoite and Schizont stages.

To test whether APA may impact overall levels of gene expression, the latter parameter was determined using the PAT-seq reads, along the lines discussed previously (Lohman et al. 2016). For the Ring vs. Schizont and Ring vs Trophozoite comparisons, the sets of genes differentially-expressed in the pairwise analyses were compared with those genes affected by APA. The results showed that 12% and 14% of the differentially-expressed genes in the Ring vs. Schizont and Ring vs Trophozoite comparisons, respectively, were also affected by APA (**Fig. 6C, Supplemental_File_S11**). APA seemed to disproportionately impact the expression of genes encoding cytoskeletal proteins, cell adhesion molecules and enzymes involved in energy metabolism and other biochemical processes (**Supplemental_File_S11**). These results suggest a modest but significant contribution of APA to stage-specific gene expression.

### Organization of m6A signatures in the context of mRNA 3’ends in *P. falciparum*

The 3’ end of mRNA is a dynamic regulatory region that integrates various signals and modifications to control gene expression. The 3’ end typically contains the last exon of the gene, which includes the coding sequence that ends with a stop codon and typically a 3’ untranslated region (3’UTR). These 3’ UTRs are known to play major roles in post-transcriptional gene regulation, including influencing mRNA localization and translation. In general, m6A marks are enriched in 3’UTRs and in the vicinity of the stop codon(Dominissini et al. 2012; Meyer et al. 2012; Lee et al. 2020). In our DART-seq based epitranscriptomic mapping of *P. falciparum* mRNAs, we found that *P. falciparum* 3’UTRs and introns contain little to no m6A marks. However, *P. falciparum* mRNA coding regions are extensively enriched with m6A marks with substantial accumulation of m6As at the CDS 3’ end (**Fig. 2B** and **Supplemental_Fig_S1**). Since accumulation of m6A residues in the last exon could impact 3’ UTR regulation(Ke et al. 2015; Luo et al. 2022), particularly affecting alternative poly(A) site choices, we first sought to measure distances of m6A sites to the 3’ UTR in the three asexual stages of *P. falciparum*. For this, random distributions of m6A sites for the Ring, Trophozoite, and Schizont stages were generated by creating 100 replicates of the DART-seq BED file for each stage, with new m6A start and end coordinates randomly assigned across the genes, while maintaining the number of random sites per gene fixed in each replicate to that observed in the original BED file. Each randomized BED file was then compared to the 3’ UTR GTF (derived from Chappell et al (Chappell et al. 2020)) to calculate the distance to the nearest m6A site from the 3’ UTR (**Fig. 7A, Supplemental_File_S12**). These density plots in **Fig. 7A** comparing the closest observed m6A site to 3’ UTR with those expected by chance across the three *P. falciparum* life stages show median observed values of 676 nt, 683 nt and 751 nt in Ring, Trophozoite and Schizont stages respectively in contrast to the randomized sites with a distance of 1222 nt, 1178 nt and 1288 nt for the above stages. Next, we determined the distance separating the nearest m6A sites identified by DART-seq in the mRNA coding region from the poly(A) sites obtained by TE-seq (**Fig. 7B**, **Supplemental_File_S13)**. In this case, the median observed values are 1487nt, 1092 nt and 1438.5 nt compared to the random values of 2434 nt, 1866 nt and 2587 nt respectively in the Ring, Trophozoite and Schizont stages. A Wilcoxon signed-rank test was performed to assess the differences in distance between observed and expected m6A sites relative to the TE-seq poly(A) tail site were significantly different, which revealed that all the three stages exhibited significant differences in distance distributions albeit Trophozoite to a lesser extent (**Fig. 7B**). **Supplemental_Fig_S8** shows an example of enrichment of a protein coding gene, Pf3D7_0406500 in which majority of the m6A sites are located at the 3’ end, primarily in the context of ‘RAC’ motif and in the vicinity of the two APA sites that are recorded by TE-seq, suggestive of the crosstalk between APA and RNA methylation.

**Figure 7.**
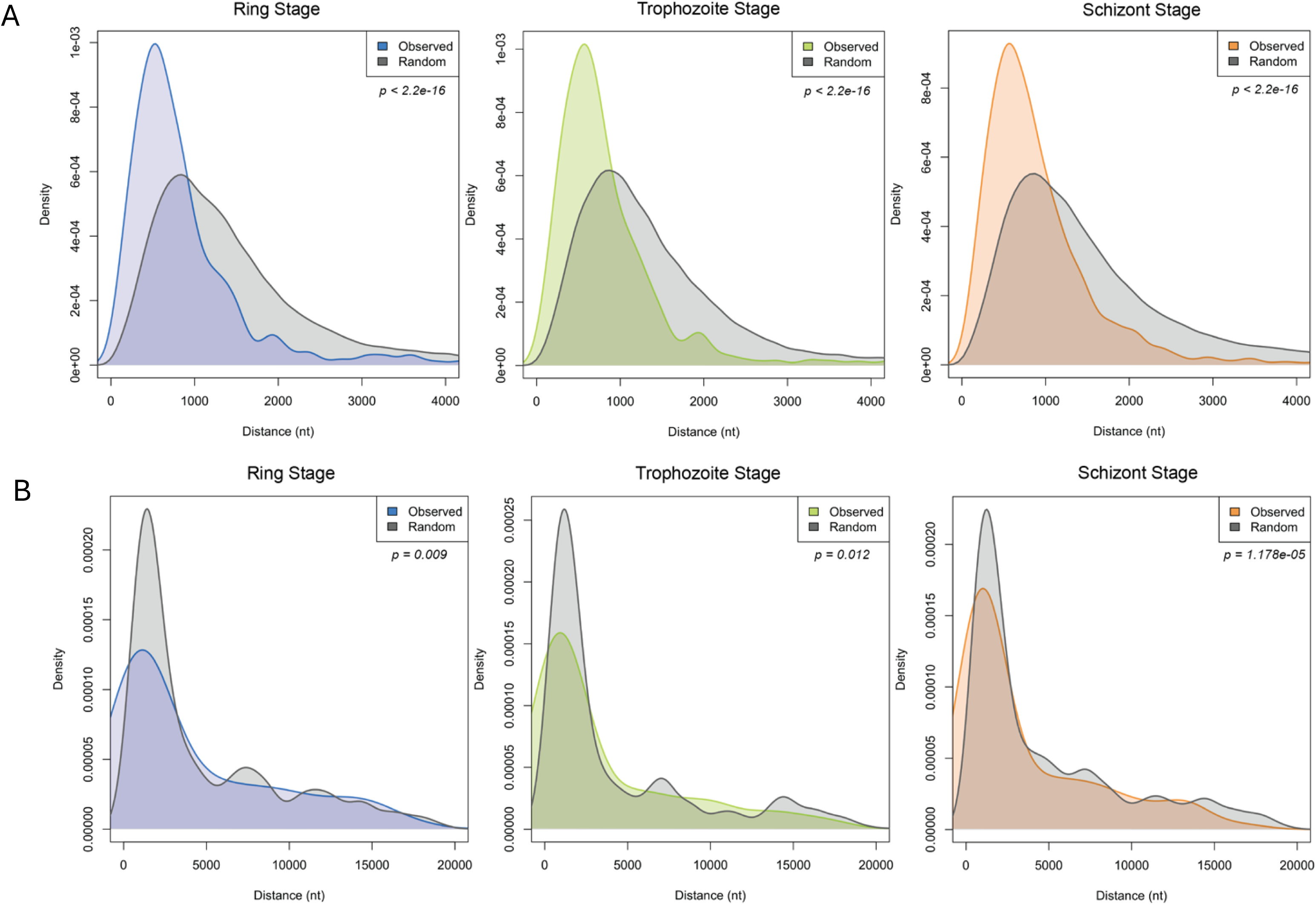
(A) Density plots comparing the closest observed m6A site to 3’ UTR with those expected by chance across the three Plasmodium falciparum life stages: Ring, Trophozoite and Schizont. A Wilcoxon signed-rank test was performed to assess the differences in distance between observed and expected m6A sites relative to the 3’ UTR. (B) Density plots comparing the closest observed m6A site to TE-seq poly(A) tail sites with those expected by chance across the three Plasmodium falciparum life stages: Ring, Trophozoite and Schizont. A Wilcoxon signed-rank test was performed to assess the differences in distance between observed and expected m6A sites relative to the TE-seq poly(A) tail site.

To explore the possible association of APA and m6A modifications, genes that showed stage-specific APA (**Supplemental_File_S11**) and that had m6A modifications (MeRIP-seq data) were assessed using an RT-qPCR assay designed to quantify the abundances of different mRNA isoforms. Specifically, to quantify cleavage abundance, we examined TE-seq PAC sites for the difference between transcript levels downstream of our chosen Poly(A) site peak (transcripts that remained uncleaved) and compared them to transcript levels upstream of our chosen peak (total transcripts) (**Supplemental_Fig_S9,** Ct values normalized relative to the control gene, H2A). We hypothesize that m6A may play a role in increased cleavage at these overlapping locations of m6A marks and PACs. To test this, we compared cleavage abundance between m6A-immunoprecipitated RNA and cellular RNA isolated by the TRIzol method. Both sets of RNA were ribosomal RNA-depleted, and DNase treated to rule out heavily modified off-target populations. We observed that, although not statistically significant, the abundance of transcripts with alternative APA sites (‘Total transcripts’) is comparatively greater compared to transcripts with canonical 3’ ends (‘Uncleaved transcripts’) for the majority of mRNAs. Further, we compared the number of APA sites from long-read RNA sequencing data (Yang et al. 2021) with m6A site abundance from DART-seq data for each gene (**Supplemental_File_S14**), however, we were not able to establish a clear connection between m6A site abundance and polyA site choice in *P. falciparum* mRNAs. Taken together, this data and m6A site preferences near the 3′ end regions of the coding sequence of *P. falciparum* mRNAs containing multiple APA sites, and significantly shorter distances between m6A sites and 3’UTRs as well as APA sites across stages suggests a possible connection of m6As to poly(A) site choices which will require further confirmatory research.

## DISCUSSION

N6-methyladenosine (m6A) is a dynamic modification that is not only crucial for regulating gene expression and RNA metabolism but also plays an essential role in the evolution of genomes(Miao et al. 2022; Shachar et al. 2024). Understanding m6A modifications provides insights into how organisms adapt, evolve, and maintain cellular functions across various environments as m6A affects the expression of genes involved in crucial biological processes, such as differentiation, proliferation, and response to stress. The malaria parasite *P. falciparum* is a member of the phylum Apicomplexa, a large group of parasitic eukaryotes that have originated early in evolution. This unicellular protozoan parasite contains about 5,300 genes in its 23-megabase nuclear genome with a mean gene length of 2.3 kb, which is longer than the average gene length in other organisms. The *P. falciparum* genome is also somewhat distinctive in its very high AT (>80% AT) content, a feature that extends to the set of Plasmodium mRNAs. In this study we have used DART-seq to map m6A landscapes in *P. falciparum* mRNAs in nucleotide-resolution and then validate the data by MeRIP-seq approach. Our results show high abundance of m6A marks in the mRNA coding regions. In mRNA, m6A methylation typically happens in a DRACH consensus motif. *P. falciparum* mRNAs on average contain five to seven m6A sites and about 15% of the m6A containing genes harbor >20 methylation marks in a single mRNA, which is similar to or slightly higher than what is recently reported in other eukaryotes (Dominissini et al. 2012; Meyer et al. 2012; Linder et al. 2015). This higher than usual m6A methylation in some genes (Supplemental File 14) could be due to the fact that *P. falciparum* genome is highly Adenosine (A)-rich and eukaryotic m6A methyltransferase complexes are known to lack specificity, methylating many adenines encountered in their recognition motifs (Zhong et al. 2008; Anderson et al. 2018; Cun et al. 2024).

Based on DART-seq and MeRIP-seq data combined, transcripts encoded by about 40% of genes in *P. falciparum* appear to be m6A methylated. The DART-seq data coverage alone accounts for about one-fourth of all genes expressed in the asexual human blood stages, including protein and RNA coding genes. Many of these sites appear to be uniquely modified in one of the three developmental stages, suggesting a possible role for m6A modifications in developmental gene regulation in malaria. In mammals, the presence of m6A modifications has been shown to enhance the degradation of mRNA(Wang et al. 2014), leading to rapid responses to environmental changes. Such adaptive mechanisms highlight the evolutionary advantage provided by m6A, allowing for swift adjustments in gene expression in response to external stimuli. The survival of *P. falciparum* in the human host is often susceptible to exposure to human immune responses, heat and oxidative stresses from drug treatments, among other stresses. Innate immune responses have been shown to contribute to the control of malaria infections, particularly human Toll-like receptor (TLR) responses that are boosted in febrile patients during natural infection with *P. falciparum*(Franklin et al. 2009; Mackowiak et al. 2021) and antimalarial drugs, such as Artemisinin, that induce *oxidative stress* (Oakley et al. 2007; Vasquez et al. 2021; Egwu et al. 2022; Vasquez et al. 2023; Pires et al. 2024). Thus, these stress conditions significantly decrease the sensitivity of the parasite to the drug as a survival response(Zhu et al. 2022; Pandit et al. 2023; Pires et al. 2023; Pires et al. 2024), but the underlying molecular mechanisms remain unknown. Since m6A’s role as a newly emerging layer of genetic control in stress responses is becoming more and more evident(Zhou et al. 2015; Qi et al. 2022; Wang et al. 2022; Ponzetti et al. 2023), it is tempting to speculate that m6A may play a regulatory role in *P. falciparum* stress responses. Interestingly, most of the heat-shock response genes (PfHSP70, PfHSP70x, PfHSP90 etc.) encode mRNAs in *P. falciparum* that are heavily methylated, supporting the above hypothesis. In addition, several genes in the malaria parasite that play key roles in nutrient uptake, antigen presentation and cytoadherence (CLAG 3.1, EXP1 and members of ETRAMP gene family, SERA and KAHRP), are m6A modified, suggesting a potential role of m6A-mediated genetic control in parasite biology.

In mammals, the motif AAUAAA and related A-rich sequences serve as polyadenylation signals (reviewed in (Boreikaitė and Passmore 2023)). These motifs are recognized by two subunits of the polyadenylation complex, CPSF30 and WDR33. In yeast, the so-called positioning element (the yeast polyadenylation signal, or yPAS) is A-rich and sits in a similar location relative to the cleavage site. The yPAS is recognized primarily by Yth1 (Boreikaitė and Passmore 2023), the yeast counterpart of CPSF30. In plants, the analogous signal is an A-rich element that is found 10-30 nts 5’ of the actual polyadenylation site. In *Arabidopsis*, this A-rich element is recognized by CPSF30 and WDR33 orthologs (Yu et al. 2019). There is evidence for a similar A-rich element in various apicomplexans 3’ regions (Stevens et al. 2018), The results presented in this report show that even within the A-rich *P. falciparum* transcriptome, there are indications of an analogous A-rich element; this is seen in the modest but clear elevated A content in 3’ ends of transcripts between 10 and 40 nts 5’ of poly(A) sites (**Fig. 5**).

The seeming ubiquity of A-rich poly(A) signals suggests that the subunits of the apicomplexan polyadenylation complex that recognize the A-rich element should be orthologous to CPSF30 and WDR33. Apicomplexans homologs of CPSF30 have been described (Stevens et al. 2018; Farhat et al. 2021; Holmes et al. 2021); these proteins resemble their plant counterparts in terms of the array of CCCH-type zinc finger motifs (three) and association with an m6A reader domain (YTH) that “reads” m6A modifications (Stevens et al. 2018; Farhat et al. 2021; Holmes et al. 2021).

The association of an m6A reader domain with the apicomplexan CPSF30 ortholog (Stevens et al. 2018; Farhat et al. 2021; Holmes et al. 2021) suggests that deposition of m6A marks near the poly(A) sites may be an additional licensing feature of polyadenylation mechanisms in *P. falciparum*. This possibility is suggested by the observation that elimination of m6A modification in *T. gondii* by mutation of the attendant methyltransferases alters mRNA polyadenylation, leading to increased transcriptional readthrough (Farhat et al. 2021; Holmes et al. 2021). Interestingly, in plants m6A modifications have been proposed to facilitate the functioning of the plant polyadenylation signal via interactions of the larger of the two plant CPSF30 isoforms (that possesses an m6A reader domain)(Hou et al. 2021; Song et al. 2021). However, the results presented in this report argue against an analogous role for m6A modifications in poly(A) site determination in *Plasmodium*, since there is no clear association of m6A modification sites with regions within 3’-UTRs that might function as poly(A) signals. This leaves unanswered the mechanism by which m6A modifications in *Plasmodium* affect poly(A) site choice and transcription termination.

Previous studies have identified individual genes in *P. falciparum* that are regulated at the level of mRNA 3’-end processing (Golightly et al. 2000; Rehkopf et al. 2000; Cann et al. 2004; Shue et al. 2004; Oguariri et al. 2006). More recently, short -read and long-read RNA sequencing technologies were used to survey genome-wide mRNA 3’ ends in *P. falciparum* (Siegel et al. 2014a; Yang et al. 2021). Importantly, the long-read RNA sequencing data identified >1500 APA sites in the intraerythrocytic stages of *P. falciparum*, and about 1/3^rd^ of the genes represented in these studies contain more than ten (10) poly(A) sites per gene (Supplemental File 14). The results presented in this report (Figs. 4 and 5) corroborate these earlier studies using two different, independent approaches; collectively, these results provide strong evidence for an involvement of alternative polyadenylation in gene expression in *P. falciparum.* Additionally, Gene Ontology (GO) enrichment analysis suggested that a significant number of genes that control mRNA metabolism, particularly, RNA transport, splicing, and polyadenylation in addition to protein metabolic processes (such as Proteosome) and redox processes are possibly regulated by m6A.

In summary, this study provides a fine-resolution map of the m6A methylation pattern in the *P. falciparum* transcriptome across asexual developmental stages of *P. falciparum* in human RBCs. This study also lays the foundation for further studies exploring the roles of m6A in polyadenylation and translation of *P. falciparum* mRNAs that are intimately involved in malaria pathogenesis and host-pathogen interactions during parasite development in intraerythrocytic stages.

## METHODS

### *Plasmodium falciparum* Culture, Synchronization and Treatments

The *P. falciparum* laboratory adapted parasite line 3D7 was cultured at 2% hematocrit in purified human erythrocytes/RBCs using standard culturing techniques (Trager and Jensen 1976) using RPMI 1640 medium (supplemented with 37.5 mM HEPES, 7 mM D-glucose, 6 mM NaOH, 25 µg of gentamicin sulfate/ml, 2 mM L-glutamine, and 10% Albumax II (Life Technologies) at a pH of 7.2 in a 5% carbon dioxide environment. The asexual blood stage parasite density was calculated routinely using thick blood films. Baseline RBC count was used to calculate the parasitaemia (parasites/µl). *P.falciparum* Infected Red Blood Cells (iRBCs) were treated with a pre-warmed aliquot of 5% D-sorbitol in RPMI at 37°C to synchronize parasite cultures in order to isolate homogeneous stages of parasites. Multiple sorbitol treatments were made to achieve >95% synchrony of parasites. Parasites were collected from three different developmental stages after stringent synchronizations: (1) Ring, 12-14 hours (2) Trophozoite, 24-26 and Schizont, 36-38 hours post-invasion. Parasites were isolated after saponin lysis, washed with incomplete medium (without Albumax) and RNA from each stage was using Trizol (Invitrogen) according to the manufacturer’s instruction and the RNA was purified as described earlier(Alvarez et al. 2021).

### Nucleotide-resolution m6A mapping with DART-seq in *P. falciparum* RBC stages

Total RNA from the three *P. falciparum* blood-stages were treated with APOBEC1-YTH and APOBEC1-YTHmut enzymes and the DART reaction was performed as described (Meyer 2019a). Briefly, two DART reactions were set up for each RNA sample - one using the wildtype (APO1-YTH) protein, the other reaction using the mutant (APO1-YTHmut) protein as control. These DART reactions were incubated at 37 °C for 4 h. After incubation, RNA from DART reactions was purified using RNeasy Micro Kit (Qiagen) according to the manufacturer’s instructions and then stored at −80 °C until downstream processing. For DART-seq, sequencing libraries were generated from 100 ng – 300ng of total RNA for each replicate using the Illumina Stranded mRNA Prep, Ligation Kit (Catalog No. 20040532) according to the manufacturer’s instructions. Sample QC was performed using a Bioanalyzer High Sensitivity DNA 1000 Chip (Agilent) and quantified using the Qubit 1X dsDNA HS Kit (Invitrogen). Paired-end sequencing was performed on an Illumina MiSeq using MiSeq Reagent Kit v2 (300-cycles). This provided a nucleotide resolution data of the m6A-modified transcriptome in *P. falciparum*.

#### Computational analyses of DART-seq data

For DART-seq based m6A mapping, sequence data from APO1-YTH (WT/m6A modified) and APO1-YTHmut (control) samples of each malaria developmental stage was input to the ‘Find_edit_site’ Perl script deployed in Bullseye tool (Zhu H), which allows detection of RNA editing sites by comparison to genomic sequence or to a control file, along with the *P. falciparum* ReFlat file. The latter is needed for identifying the m6A editing sites only in regions specified in an annotation file. This allows annotation of m6A sites to a specified set of genomic features listed in the ReFlat file. ReFlat files can be downloaded from UCSC table browser, via the reFlat table of the NCBI RefSeq track or a custom GTF annotation file can be converted to a reFlat using the provided ‘gtf2genepresd perl script’ implemented in Bullseye tool source code. The output of running the ‘Find_edit_site’ perl script was a bed file consisting of all m6A mRNA modified sites for each *Plasmodium falciparum* RBC stage. At this stage, two matrix files of DART-seq data corresponding to m6A modified and control samples data for each stage were created and analyzed.

#### MeRIP-Seq with *P. falciparum* RBC stages

To validate DART-seq peaks, two replicates of MeRIP-Seq dataset were created using a modified MeRIP-seq (Meyer et al. 2012) method. In this method, mixed -stage RNAs from asynchronous *P. falciparum* cultures were isolated from asynchronous *in vitro* cultures and treated with Turbo DNAse (Thermo Fisher) according to manufacturer’s instructions. Following DNAse treatment, Ribosomal RNAs (rRNAs) were removed using a custom assay developed specifically for *P. falciparum* rRNAs (Alvarez et al. 2021). Following this, immunoprecipitation of m6A -enriched mRNAs were performed using a rabbit monoclonal antibody specific for N6-Methyladenosine (m6A) (New England Biolabs, Catalog No. E1611) and then purified with Protein G Magnetic Beads (New England Biolabs, Catalog No. S1430). RNA clean-up and concentrations were accomplished using Monarch RNA Cleanup Kit (10 μg) (NEB #T2030). RNA Library prep was done using the Illumina Stranded mRNA Prep, Ligation Kit (Catalog No. 20040532) according to the manufacturer’s instructions followed by Illumina Sequencing on an Illumina Miseq platform.

Reads underwent quality assessment using FASTQC and were subsequently trimmed using TrimGalore (Krueger 2015). Alignment to the *P. falciparum* genome (GCF_000002765.6) was performed with HISAT2 (Lee et al. 2020). SAMTOOLS was employed to generate BAM files, which served as input for peak calling using Macs2 callpeak with the -broadpeaks parameter. This process yielded 4006 and 4099 peaks for replicate 1 and replicate 2, respectively. Consensus peaks, totaling 3,506, were determined by intersecting peaks identified in the two replicates using bedtools (Quinlan and Hall 2010). The resulting consensus peaks were utilized to validate the presence of DART-seq hits in the MeRIP-seq data.

#### Motif Discovery

For the identification of m6A motifs within the DART-seq sites across the three Plasmodium stages, an 11-bp sequence centered on the m6A site was created by extending the coordinates 5 bp to the left and right of the m6A site. Bedtools slop was employed for coordinate extension, followed by the extraction of the corresponding sequences using bedtools getfasta. Subsequently, motif discovery was conducted using MEME (Bailey et al. 2015).

### PAT-seq and TE-seq: library preparation and deep sequencing of RNA 3’ ends

Two methods were used to prepare libraries for genome-wide characterization of poly(A) sites in *P. falciparum*. One method (termed in this report as PAT-seq) is that described elsewhere (Wu et al. 2011; Ma et al. 2014; Pati et al. 2015) and entails (sequentially) RNA fragmentation, purification of poly(A) containing RNAs using oligo-dT beads, and cDNA synthesis using oligo-dT-based priming. The second method (abbreviated hereafter as TE-seq, for ‘Three prime/3’ End Seq’) is more painstaking and is intended to profile RNA 3’ ends without RT priming using oligo-dT. For this, a protocol described in detail elsewhere (Chakrabarti et al. 2020) was modified. An RNA adaptor (R1-RNA adaptor, **Supplemental_File_S1**) was ligated to the 3’ ends of total RNA (1 µg) using T4 RNA ligase I (ssRNA ligase; New England Biolabs Catalog M0204) as per the manufacturer’s recommendations. Ligation products were recovered into 30 µL RNAse-free water using Qiagen MinElute RNA cleanup kits (Cat. 74204). Adaptor-ligated RNA was then treated with RNAse H and an oligonucleotide cocktail (**Supplemental_File_S1**) designed to remove *P. falciparum* ribosomal RNAs. 100 µL reactions contained the entire ligated RNA sample, 500 pmol of each of the oligonucleotides, 10 units of thermostable RNAse H (Epicentre), all in the supplier’s reaction buffer. After 1 hour at 60°C, RNAse H-treated ligated RNAs were recovered into 15 µL using Qiagen RNA Cleanup kits and fragmented by mixing the RNA sample (in a volume of 14 µL) with 1 µL (100 pmol) of the appropriate RT primer (see **Supplemental_File_S1**) and 5 µL 10X SMARTScribe First Strand Buffer (Clontech) and heating to 95°C for exactly 2 minutes. Reverse transcription reactions were then initiated by the addition of dNTPs (2.5 µL of a mixture of 10 mM of each dNTP), DTT (1 µL 20 mM DTT), and SMARTScribe (1 µL). After 60 minutes at 42°C, 1 µL of a strand-switching oligonucleotide (SMART7.5, **Supplemental_File_S1**) and 1 µL of SMARTscribe was added and reactions incubated at 42°C for an additional 60 minutes. Reactions were stopped by heating to 70°C for 5 minutes. Following reverse transcription and strand-switching, cDNAs were purified twice using SPRI beads (MagBio Genomics Inc. Catalog AC-60050); for this, 16.25 µL of the bead suspension was added to the cDNA reaction, the samples were mixed thoroughly, and the suspension incubated at room temperature for 8 minutes. Beads were collected using a magnet stand (NEB S1506S), washed three times with 100 µL 80% ethanol, air-dried for 5 minutes, and cDNAs eluted into 25 µL of RNAse-free water. Purified cDNAs were PCR amplified using one primer from the R4-RT series and PE-PCR2 (**Supplemental_File_S1**); the R4-RT primers possess Illumina-specific sequences at their 5’ ends, followed by a 5 nt region containing sample-specific bar codes, and then by sequences complementary to the R5-RT Master primer. PCR products were size selected using agarose gel electrophoresis and amplified again the PE-PCR1 and PE-PCR2 primers (**Supplemental_File_S1**). Final PCR products were purified using one round of SPRI bead as described above. Qualities of the final libraries were assessed using agarose gel electrophoresis and concentrations were measured with Qubit fluorometer with Qubit® dsDNA HS assay kit (Life Technologies). Statistics concerning the sequencing data (read numbers, mapping statistics, etc.) are provided in **Supplemental_File_S2**.

PAT-seq and TE-seq reads were analyzed as described in detail in **Supplemental_File_S3.** Briefly, raw sequencing reads were demultiplexed, trimmed to remove the 5’-oligo-dT tracts and sequencing adapters, and mapped to the *Plasmodium falciparum* genome (Gardner et al. 2002). Mapped reads were then trimmed to a length of one nt, preserving genome mapping coordinates. Trimmed mapped reads were used to assess poly(A) site location and usage. Trimmed mapped reads were also used to determine overall gene expression. For the plot shown in Fig. 5C, genomic locations of motifs with the general sequence N_6_BA_6_ were collected, extended by 100 nts in both directions, and the nucleotide compositions plotted as shown.

### Real-time Quantitative PCR (RT-qPCR) assays to detect Alternative Polyadenylation (APA)

Total RNAs extracted from *Plasmodium falciparum* life stages (see above) were used for RT-qPCR analysis. Total RNA was reverse transcribed into cDNA with Superscript II (Invitrogen). Reverse transcription was performed at 25 °C for 10 min, followed at 42 °C for 50 min and then inactivation at 70 °C for 15 min. The cDNA samples were diluted and stored at 4 °C. Quantitative real-time PCR (RT-qPCR) analysis was performed with PowerUp™ SYBR™ Green Master Mix for qPCR (Applied Biosystems, ABI) on the ABI QuantStudio 3 system. Primer sequences used in these assays are in the **Supplemental_File_S4**.

## DATA ACCESS

The m6A sequencing data (DART-seq), MeRIP-seq and mRNA 3’-end sequencing (TE-seq and PAT-seq) data generated in this study have been submitted to the NCBI BioProject database (https://www.ncbi.nlm.nih.gov/bioproject/) under accession number PRJNA1176112

## AUTHOR DECLARATION

All authors have read and approved the current version of this manuscript and have agreed to its submission.

## ACKNOWLEDGEMENTS

We thank Bo Zheng for help in culturing *P. falciparum* asexual stages, Karen Lopez for help with Illumina RNA-seq experiments and Shaoyu Li at the UNC Charlotte Department of Math and Statistics for discussions.

## REFERENCES

Alvarez DR, Ospina A, Barwell T, Zheng B, Dey A, Li C, Basu S, Shi X, Kadri S, Chakrabarti K. 2021. The RNA Structurome in the Asexual Blood Stages of Malaria Pathogen Plasmodium falciparum. RNA Biol doi:10.1080/15476286.2021.1926747.

Anderson SJ, Kramer MC, Gosai SJ, Yu X, Vandivier LE, Nelson ADL, Anderson ZD, Beilstein MA, Fray RG, Lyons E et al. 2018. N(6)-Methyladenosine Inhibits Local Ribonucleolytic Cleavage to Stabilize mRNAs in Arabidopsis. Cell Rep 25: 1146–1157 e1143.

Bailey TL, Johnson J, Grant CE, Noble WS. 2015. The MEME Suite. Nucleic Acids Res 43: W39–49.

Balacco DL, Soller M. 2019. The m(6)A Writer: Rise of a Machine for Growing Tasks. Biochemistry 58: 363–378.

Bartosovic M, Molares HC, Gregorova P, Hrossova D, Kudla G, Vanacova S. 2017. N6-methyladenosine demethylase FTO targets pre-mRNAs and regulates alternative splicing and 3’-end processing. Nucleic Acids Res 45: 11356–11370.

Baumgarten S, Bryant JM, Sinha A, Reyser T, Preiser PR, Dedon PC, Scherf A. 2019. Transcriptome-wide dynamics of extensive m(6)A mRNA methylation during Plasmodium falciparum blood-stage development. Nat Microbiol 4: 2246–2259.

Beeson JG, Drew DR, Boyle MJ, Feng G, Fowkes FJ, Richards JS. 2016. Merozoite surface proteins in red blood cell invasion, immunity and vaccines against malaria. FEMS Microbiol Rev 40: 343–372.

Bell SA, Shen C, Brown A, Hunt AG. 2016. Experimental Genome-Wide Determination of RNA Polyadenylation in Chlamydomonas reinhardtii. PLoS One 11: e0146107.

Boreikaitė V, Passmore LA. 2023. 3’-End Processing of Eukaryotic mRNA: Machinery, Regulation, and Impact on Gene Expression. Annual review of biochemistry 92: 199–225.

Campbell TL, De Silva EK, Olszewski KL, Elemento O, Llinas M. 2010. Identification and genome-wide prediction of DNA binding specificities for the ApiAP2 family of regulators from the malaria parasite. PLoS Pathog 6: e1001165.

Cann H, Brown SV, Oguariri RM, Golightly LM. 2004. 3’ UTR signals necessary for expression of the Plasmodium gallinaceum ookinete protein, Pgs28, share similarities with those of yeast and plants. Mol Biochem Parasitol 137: 239–245.

Catacalos C, Krohannon A, Somalraju S, Meyer KD, Janga SC, Chakrabarti K. 2022. Epitranscriptomics in parasitic protists: Role of RNA chemical modifications in posttranscriptional gene regulation. PLoS Pathog 18: e1010972.

Chakrabarti M, de Lorenzo L, Abdel-Ghany SE, Reddy ASN, Hunt AG. 2020. Wide-ranging transcriptome remodelling mediated by alternative polyadenylation in response to abiotic stresses in Sorghum. Plant J 102: 916–930.

Chakrabarti M, Hunt AG. 2015. CPSF30 at the Interface of Alternative Polyadenylation and Cellular Signaling in Plants. Biomolecules 5: 1151–1168.

Chan S, Choi EA, Shi Y. 2011. Pre-mRNA 3’-end processing complex assembly and function. Wiley Interdiscip Rev RNA 2: 321–335.

Chappell L, Ross P, Orchard L, Russell TJ, Otto TD, Berriman M, Rayner JC, Llinas M. 2020. Refining the transcriptome of the human malaria parasite Plasmodium falciparum using amplification-free RNA-seq. BMC Genomics 21: 395.

Chen L, Fu Y, Hu Z, Deng K, Song Z, Liu S, Li M, Ou X, Wu R, Liu M et al. 2022. Nuclear m(6) A reader YTHDC1 suppresses proximal alternative polyadenylation sites by interfering with the 3’ processing machinery. EMBO Rep 23: e54686.

Collins CR, Hackett F, Strath M, Penzo M, Withers-Martinez C, Baker DA, Blackman MJ. 2013. Malaria parasite cGMP-dependent protein kinase regulates blood stage merozoite secretory organelle discharge and egress. PLoS Pathog 9: e1003344.

Cun Y, Guo W, Ma B, Okuno Y, Wang J. 2024. Decoding the specificity of m(6)A RNA methylation and its implication in cancer therapy. Mol Ther 32: 2461–2469.

de Lorenzo L, Sorenson R, Bailey-Serres J, Hunt AG. 2017. Noncanonical Alternative Polyadenylation Contributes to Gene Regulation in Response to Hypoxia. Plant Cell 29: 1262–1277.

Desai SA, Krogstad DJ, McCleskey EW. 1993. A nutrient-permeable channel on the intraerythrocytic malaria parasite. Nature 362: 643–646.

Di Giammartino DC, Nishida K, Manley JL. 2011. Mechanisms and consequences of alternative polyadenylation. Mol Cell 43: 853–866.

Dominissini D, Moshitch-Moshkovitz S, Schwartz S, Salmon-Divon M, Ungar L, Osenberg S, Cesarkas K, Jacob-Hirsch J, Amariglio N, Kupiec M et al. 2012. Topology of the human and mouse m6A RNA methylomes revealed by m6A-seq. Nature 485: 201–206.

Dutta T, Singh H, Edkins AL, Blatch GL. 2022. Hsp90 and Associated Co-Chaperones of the Malaria Parasite. Biomolecules 12.

Egwu CO, Perio P, Augereau JM, Tsamesidis I, Benoit-Vical F, Reybier K. 2022. Resistance to artemisinin in falciparum malaria parasites: A redox-mediated phenomenon. Free Radic Biol Med 179: 317–327.

Elkon R, Ugalde AP, Agami R. 2013. Alternative cleavage and polyadenylation: extent, regulation and function. Nat Rev Genet 14: 496–506.

Farhat DC, Bowler MW, Communie G, Pontier D, Belmudes L, Mas C, Corrao C, Coute Y, Bougdour A, Lagrange T et al. 2021. A plant-like mechanism coupling m6A reading to polyadenylation safeguards transcriptome integrity and developmental gene partitioning in Toxoplasma. Elife 10.

Franklin BS, Parroche P, Ataide MA, Lauw F, Ropert C, de Oliveira RB, Pereira D, Tada MS, Nogueira P, da Silva LH et al. 2009. Malaria primes the innate immune response due to interferon-gamma induced enhancement of toll-like receptor expression and function. Proc Natl Acad Sci U S A 106: 5789–5794.

G. Bindea BM, H. Hackl, P. Charoentong, M. Tosolini, A. Kirilovsky, W.-, H. Fridman FPe, Z. Trajanoski, J. Galon. ClueGO: a Cytoscape plug-in to decipher functionally grouped gene ontology and pathway annotation networks, Bioinformatics (Oxford, England) 25 (8) (2009) 1091–1093, 10.1093/bioinformatics/btp101.

Gardner MJ, Hall N, Fung E, White O, Berriman M, Hyman RW, Carlton JM, Pain A, Nelson KE, Bowman S et al. 2002. Genome sequence of the human malaria parasite Plasmodium falciparum. Nature 419: 498–511.

Goldberg DE, Zimmerberg J. 2020. Hardly Vacuous: The Parasitophorous Vacuolar Membrane of Malaria Parasites. Trends Parasitol 36: 138–146.

Golightly LM, Mbacham W, Daily J, Wirth DF. 2000. 3’ UTR elements enhance expression of Pgs28, an ookinete protein of Plasmodium gallinaceum. Mol Biochem Parasitol 105: 61–70.

Govindaraju G, Kadumuri RV, Sethumadhavan DV, Jabeena CA, Chavali S, Rajavelu A. 2020. N(6)-Adenosine methylation on mRNA is recognized by YTH2 domain protein of human malaria parasite Plasmodium falciparum. Epigenetics Chromatin 13: 33.

Grover M, Chaubey S, Ranade S, Tatu U. 2013. Identification of an exported heat shock protein 70 in Plasmodium falciparum. Parasite 20: 2.

Hollin T, Le Roch KG. 2020. From Genes to Transcripts, a Tightly Regulated Journey in Plasmodium. Front Cell Infect Microbiol 10: 618454.

Holmes MJ, Padgett LR, Bastos MS, Sullivan WJ, Jr. 2021. m6A RNA methylation facilitates pre-mRNA 3’-end formation and is essential for viability of Toxoplasma gondii. PLoS Pathog 17: e1009335.

Hou Y, Sun J, Wu B, Gao Y, Nie H, Nie Z, Quan S, Wang Y, Cao X, Li S. 2021. CPSF30-L-mediated recognition of mRNA m(6)A modification controls alternative polyadenylation of nitrate signaling-related gene transcripts in Arabidopsis. Mol Plant 14: 688–699.

Hunt AG, Howe DK, Brown A, Yeargan M. 2021. Transcriptional dynamics in the protozoan parasite Sarcocystis neurona and mammalian host cells after treatment with a specific inhibitor of apicomplexan mRNA polyadenylation. PLoS One 16: e0259109.

Kasowitz SD, Ma J, Anderson SJ, Leu NA, Xu Y, Gregory BD, Schultz RM, Wang PJ. 2018. Nuclear m6A reader YTHDC1 regulates alternative polyadenylation and splicing during mouse oocyte development. PLoS Genet 14: e1007412.

Ke S, Alemu EA, Mertens C, Gantman EC, Fak JJ, Mele A, Haripal B, Zucker-Scharff I, Moore MJ, Park CY et al. 2015. A majority of m6A residues are in the last exons, allowing the potential for 3’ UTR regulation. Genes Dev 29: 2037–2053.

Krueger F. 2015. A wrapper tool around Cutadapt and FastQC to consistently apply quality and adapter trimming to FastQ files. Vol 516, p. 517.

Lee Y, Choe J, Park OH, Kim YK. 2020. Molecular Mechanisms Driving mRNA Degradation by m(6)A Modification. Trends Genet 36: 177–188.

Li XQ, Du D. 2014. Motif types, motif locations and base composition patterns around the RNA polyadenylation site in microorganisms, plants and animals. BMC Evol Biol 14: 162.

Liao S, Sun H, Xu C. 2018. YTH Domain: A Family of N(6)-methyladenosine (m(6)A) Readers. Genomics Proteomics Bioinformatics 16: 99–107.

Linder B, Grozhik AV, Olarerin-George AO, Meydan C, Mason CE, Jaffrey SR. 2015. Single-nucleotide-resolution mapping of m6A and m6Am throughout the transcriptome. Nat Methods 12: 767–772.

Liu L, Zeng S, Jiang H, Zhang Y, Guo X, Wang Y. 2019. Differential m6A methylomes between two major life stages allows potential regulations in Trypanosoma brucei. Biochem Biophys Res Commun 508: 1286–1290.

Llinas M, Deitsch KW, Voss TS. 2008. Plasmodium gene regulation: far more to factor in. Trends Parasitol 24: 551–556.

Lohman BK, Weber JN, Bolnick DI. 2016. Evaluation of TagSeq, a reliable low-cost alternative for RNAseq. Mol Ecol Resour 16: 1315–1321.

Luo Z, Zhang J, Fei J, Ke S. 2022. Deep learning modeling m(6)A deposition reveals the importance of downstream cis-element sequences. Nat Commun 13: 2720.

Lutz CS, Moreira A. 2011. Alternative mRNA polyadenylation in eukaryotes: an effective regulator of gene expression. WIREs RNA 2: 23–31.

Ma L, Pati PK, Liu M, Li QQ, Hunt AG. 2014. High throughput characterizations of poly(A) site choice in plants. Methods 67: 74–83.

Mackowiak PA, Chervenak FA, Grunebaum A. 2021. Defining Fever. Open Forum Infect Dis 8: ofab161.

Maier AG, Cooke BM, Cowman AF, Tilley L. 2009. Malaria parasite proteins that remodel the host erythrocyte. Nat Rev Microbiol 7: 341–354.

Mair GR, Braks JA, Garver LS, Wiegant JC, Hall N, Dirks RW, Khan SM, Dimopoulos G, Janse CJ, Waters AP. 2006. Regulation of sexual development of Plasmodium by translational repression. Science 313: 667–669.

Mao Y, Dong L, Liu XM, Guo J, Ma H, Shen B, Qian SB. 2019. m(6)A in mRNA coding regions promotes translation via the RNA helicase-containing YTHDC2. Nat Commun 10: 5332.

Mayr C. 2016. Evolution and Biological Roles of Alternative 3’UTRs. Trends Cell Biol 26: 227–237.

Mesen-Ramirez P, Bergmann B, Tran TT, Garten M, Stacker J, Naranjo-Prado I, Hohn K, Zimmerberg J, Spielmann T. 2019. EXP1 is critical for nutrient uptake across the parasitophorous vacuole membrane of malaria parasites. PLoS Biol 17: e3000473.

Meyer KD. 2019a. DART-seq: an antibody-free method for global m(6)A detection. Nat Methods 16: 1275–1280.

Meyer KD. 2019b. m(6)A-mediated translation regulation. Biochim Biophys Acta Gene Regul Mech 1862: 301–309.

Meyer KD, Jaffrey SR. 2014. The dynamic epitranscriptome: N6-methyladenosine and gene expression control. Nat Rev Mol Cell Biol 15: 313–326.

Meyer KD, Jaffrey SR. 2017. Rethinking m(6)A Readers, Writers, and Erasers. Annu Rev Cell Dev Biol 33: 319–342.

Meyer KD, Patil DP, Zhou J, Zinoviev A, Skabkin MA, Elemento O, Pestova TV, Qian SB, Jaffrey SR. 2015. 5’ UTR m(6)A Promotes Cap-Independent Translation. Cell 163: 999–1010.

Meyer KD, Saletore Y, Zumbo P, Elemento O, Mason CE, Jaffrey SR. 2012. Comprehensive analysis of mRNA methylation reveals enrichment in 3’ UTRs and near stop codons. Cell 149: 1635–1646.

Miao Z, Zhang T, Xie B, Qi Y, Ma C. 2022. Evolutionary Implications of the RNA N6-Methyladenosine Methylome in Plants. Mol Biol Evol 39.

Molinie B, Wang J, Lim KS, Hillebrand R, Lu ZX, Van Wittenberghe N, Howard BD, Daneshvar K, Mullen AC, Dedon P et al. 2016. m(6)A-LAIC-seq reveals the census and complexity of the m(6)A epitranscriptome. Nat Methods 13: 692–698.

Oakley MS, Kumar S, Anantharaman V, Zheng H, Mahajan B, Haynes JD, Moch JK, Fairhurst R, McCutchan TF, Aravind L. 2007. Molecular factors and biochemical pathways induced by febrile temperature in intraerythrocytic Plasmodium falciparum parasites. Infect Immun 75: 2012–2025.

Oguariri RM, Dunn JM, Golightly LM. 2006. 3’ gene regulatory elements required for expression of the Plasmodiumfalciparum developmental protein, Pfs25. Mol Biochem Parasitol 146: 163–172.

Pandit K, Surolia N, Bhattacharjee S, Karmodiya K. 2023. The many paths to artemisinin resistance in Plasmodium falciparum. Trends Parasitol 39: 1060–1073.

Pati PK, Ma L, Hunt AG. 2015. Genome-wide determination of poly(A) site choice in plants. Methods Mol Biol 1255: 159–174.

Pires CV, Cassandra D, Xu S, Laleu B, Burrows JN, Adams JH. 2024. Oxidative stress changes the effectiveness of artemisinin in Plasmodium falciparum. mBio 15: e0316923.

Pires CV, Chawla J, Simmons C, Gibbons J, Adams JH. 2023. Heat-shock responses: systemic and essential ways of malaria parasite survival. Curr Opin Microbiol 73: 102322.

Ponzetti M, Rucci N, Falone S. 2023. RNA methylation and cellular response to oxidative stress-promoting anticancer agents. Cell Cycle 22: 870–905.

Proudfoot NJ. 2011. Ending the message: poly(A) signals then and now. Genes Dev 25: 1770–1782.

Qi Y, Zhang Y, Zhang J, Wang J, Li Q. 2022. The alteration of N6-methyladenosine (m6A) modification at the transcriptome-wide level in response of heat stress in bovine mammary epithelial cells. BMC Genomics 23: 829.

Quinlan AR, Hall IM. 2010. BEDTools: a flexible suite of utilities for comparing genomic features. Bioinformatics 26: 841–842.

Rehkopf DH, Gillespie DE, Harrell MI, Feagin JE. 2000. Transcriptional mapping and RNA processing of the Plasmodium falciparum mitochondrial mRNAs. Mol Biochem Parasitol 105: 91–103.

Shachar R, Dierks D, Garcia-Campos MA, Uzonyi A, Toth U, Rossmanith W, Schwartz S. 2024. Dissecting the sequence and structural determinants guiding m6A deposition and evolution via inter- and intra-species hybrids. Genome Biol 25: 48.

Shue P, Brown SV, Cann H, Singer EF, Appleby S, Golightly LM. 2004. The 3’ UTR elements of P. gallinaceum protein Pgs28 are functionally distinct from those of human cells. Mol Biochem Parasitol 137: 355–359.

Siegel TN, Hon CC, Zhang Q, Lopez-Rubio JJ, Scheidig-Benatar C, Martins RM, Sismeiro O, Coppee JY, Scherf A. 2014a. Strand-specific RNA-Seq reveals widespread and developmentally regulated transcription of natural antisense transcripts in Plasmodium falciparum. BMC Genomics 15: 150.

Siegel TN, Hon CC, Zhang Q, Lopez-Rubio JJ, Scheidig-Benatar C, Martins RM, Sismeiro O, Coppee JY, Scherf A. 2014b. Strand-specific RNA-Seq reveals widespread and developmentally regulated transcription of natural antisense transcripts in Plasmodium falciparum. BMC Genomics 15: 150.

Sinha A, Baumgarten S, Distiller A, McHugh E, Chen P, Singh M, Bryant JM, Liang J, Cecere G, Dedon PC et al. 2021. Functional Characterization of the m(6)A-Dependent Translational Modulator PfYTH.2 in the Human Malaria Parasite. mBio 12.

Song P, Yang J, Wang C, Lu Q, Shi L, Tayier S, Jia G. 2021. Arabidopsis N(6)-methyladenosine reader CPSF30-L recognizes FUE signals to control polyadenylation site choice in liquid-like nuclear bodies. Mol Plant 14: 571–587.

Spielmann T, Gardiner DL, Beck HP, Trenholme KR, Kemp DJ. 2006. Organization of ETRAMPs and EXP-1 at the parasite-host cell interface of malaria parasites. Mol Microbiol 59: 779–794.

Stevens AT, Howe DK, Hunt AG. 2018. Characterization of mRNA polyadenylation in the apicomplexa. PLoS One 13: e0203317.

Tegowski M, Flamand MN, Meyer KD. 2022. scDART-seq reveals distinct m(6)A signatures and mRNA methylation heterogeneity in single cells. Mol Cell doi:10.1016/j.molcel.2021.12.038.

Tian B, Hu J, Zhang H, Lutz CS. 2005. A large-scale analysis of mRNA polyadenylation of human and mouse genes. Nucleic Acids Res 33: 201–212.

Tian B, Manley JL. 2016. Alternative polyadenylation of mRNA precursors. Nat Rev Mol Cell Biol doi:10.1038/nrm.2016.116.

Tian B, Pan Z, Lee JY. 2007. Widespread mRNA polyadenylation events in introns indicate dynamic interplay between polyadenylation and splicing. Genome Res 17: 156–165.

Tilley L, Sougrat R, Lithgow T, Hanssen E. 2008. The twists and turns of Maurer’s cleft trafficking in P. falciparum-infected erythrocytes. Traffic 9: 187–197.

Tinto-Font E, Cortes A. 2022. Malaria parasites do respond to heat. Trends Parasitol 38: 435–449.

Tinto-Font E, Michel-Todo L, Russell TJ, Casas-Vila N, Conway DJ, Bozdech Z, Llinas M, Cortes A. 2021. A heat-shock response regulated by the PfAP2-HS transcription factor protects human malaria parasites from febrile temperatures. Nat Microbiol 6: 1163–1174.

Trager W, Jensen JB. 1976. Human malaria parasites in continuous culture. Science 193: 673–675.

Vasquez M, Sica M, Namazzi R, Opoka RO, Sherman J, Datta D, Duran-Frigola M, Ssenkusu JM, John CC, Conroy AL et al. 2023. Xanthine oxidase levels and immune dysregulation are independently associated with anemia in Plasmodium falciparum malaria. Sci Rep 13: 14720.

Vasquez M, Zuniga M, Rodriguez A. 2021. Oxidative Stress and Pathogenesis in Malaria. Front Cell Infect Microbiol 11: 768182.

Viegas IJ, de Macedo JP, Serra L, De Niz M, Temporao A, Silva Pereira S, Mirza AH, Bergstrom E, Rodrigues JA, Aresta-Branco F et al. 2022. N(6)-methyladenosine in poly(A) tails stabilize VSG transcripts. Nature 604: 362–370.

Waller KL, Cooke BM, Nunomura W, Mohandas N, Coppel RL. 1999. Mapping the binding domains involved in the interaction between the Plasmodium falciparum knob-associated histidine-rich protein (KAHRP) and the cytoadherence ligand P. falciparum erythrocyte membrane protein 1 (PfEMP1). J Biol Chem 274: 23808–23813.

Wang L, Zhan G, Maimaitiyiming Y, Su Y, Lin S, Liu J, Su K, Lin J, Shen S, He W et al. 2022. m(6)A modification confers thermal vulnerability to HPV E7 oncotranscripts via reverse regulation of its reader protein IGF2BP1 upon heat stress. Cell Rep 41: 111546.

Wang X, Lu Z, Gomez A, Hon GC, Yue Y, Han D, Fu Y, Parisien M, Dai Q, Jia G et al. 2014. N6-methyladenosine-dependent regulation of messenger RNA stability. Nature 505: 117–120.

Wang X, Zhao BS, Roundtree IA, Lu Z, Han D, Ma H, Weng X, Chen K, Shi H, He C. 2015. N(6)-methyladenosine Modulates Messenger RNA Translation Efficiency. Cell 161: 1388–1399.

Wu X, Gaffney B, Hunt AG, Li QQ. 2014. Genome-wide determination of poly(A) sites in Medicago truncatula: evolutionary conservation of alternative poly(A) site choice. BMC Genomics 15: 615.

Wu X, Liu M, Downie B, Liang C, Ji G, Li QQ, Hunt AG. 2011. Genome-wide landscape of polyadenylation in Arabidopsis provides evidence for extensive alternative polyadenylation. Proc Natl Acad Sci U S A 108: 12533–12538.

Xing D, Li QQ. 2011. Alternative polyadenylation and gene expression regulation in plants. Wiley Interdiscip Rev RNA 2: 445–458.

Yang M, Shang X, Zhou Y, Wang C, Wei G, Tang J, Zhang M, Liu Y, Cao J, Zhang Q. 2021. Full-Length Transcriptome Analysis of Plasmodium falciparum by Single-Molecule Long-Read Sequencing. Front Cell Infect Microbiol 11: 631545.

Yu Z, Lin J, Li QQ. 2019. Transcriptome Analyses of FY Mutants Reveal Its Role in mRNA Alternative Polyadenylation. Plant Cell 31: 2332–2352.

Yue Y, Liu J, Cui X, Cao J, Luo G, Zhang Z, Cheng T, Gao M, Shu X, Ma H et al. 2018. VIRMA mediates preferential m(6)A mRNA methylation in 3’UTR and near stop codon and associates with alternative polyadenylation. Cell Discov 4: 10.

Zhang M, Zhai Y, Zhang S, Dai X, Li Z. 2020. Roles of N6-Methyladenosine (m(6)A) in Stem Cell Fate Decisions and Early Embryonic Development in Mammals. Front Cell Dev Biol 8: 782.

Zhong S, Li H, Bodi Z, Button J, Vespa L, Herzog M, Fray RG. 2008. MTA is an Arabidopsis messenger RNA adenosine methylase and interacts with a homolog of a sex-specific splicing factor. Plant Cell 20: 1278–1288.

Zhou J, Wan J, Gao X, Zhang X, Jaffrey SR, Qian SB. 2015. Dynamic m(6)A mRNA methylation directs translational control of heat shock response. Nature 526: 591–594.

Zhu H YX, Holley CL, Meyer KD. Improved Methods for Deamination-Based m6A Detection. . Front Cell Dev Biol. 2022 Apr 27;10:888279. doi: 10.3389/fcell.2022.888279. PMID: 35573664; PMCID: PMC9092492.

Zhu L, van der Pluijm RW, Kucharski M, Nayak S, Tripathi J, White NJ, Day NPJ, Faiz A, Phyo AP, Amaratunga C et al. 2022. Artemisinin resistance in the malaria parasite, Plasmodium falciparum, originates from its initial transcriptional response. Commun Biol 5: 274.

