## Supplemental File for "Nucleotide-resolution Mapping of RNA N6-Methyladenosine (m6A) modifications and comprehensive analysis of global polyadenylation events in mRNA 3’ end processing in malaria pathogen *Plasmodium falciparum*"

#### SUPPLEMENTARY FIGURE LEGENDS

**Supplementary Figure 1.** Density plots representing m6A modifications across normalized gene length for the three falciparum life stages. Kruskal-Wallis test was conducted to assess the differences in m6A distribution across the three life stages.

**Supplementary Figure 2.** ClueGo enrichment network illustration biological pathways associated with the 835 unique m6A-modified genes in the Ring stage.

**Supplementary Figure 3.** ClueGo enrichment network illustration biological pathways associated with the 713 unique m6A-modified genes in the Trophozoite stage.

**Supplementary Figure 4.** ClueGo enrichment network illustration biological pathways associated with the 1713 unique m6A-modified genes in the Schizont stage.

**Supplementary Figure 5.** Workflow for TE-seq experiment to determine PolyA sites in Plasmodium falciparum using a 3'end sequencing strategy.

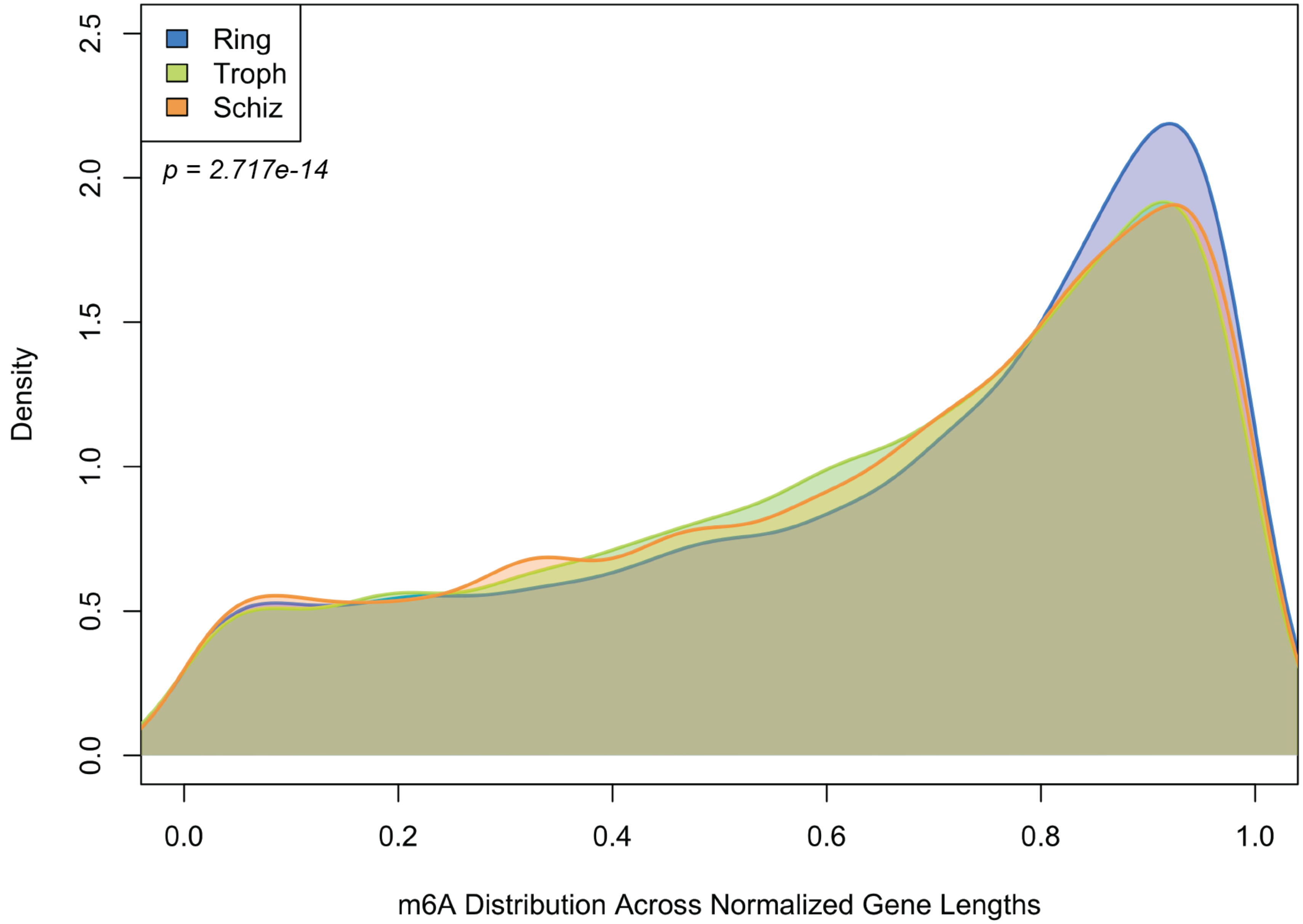


Supplementary Figure-1

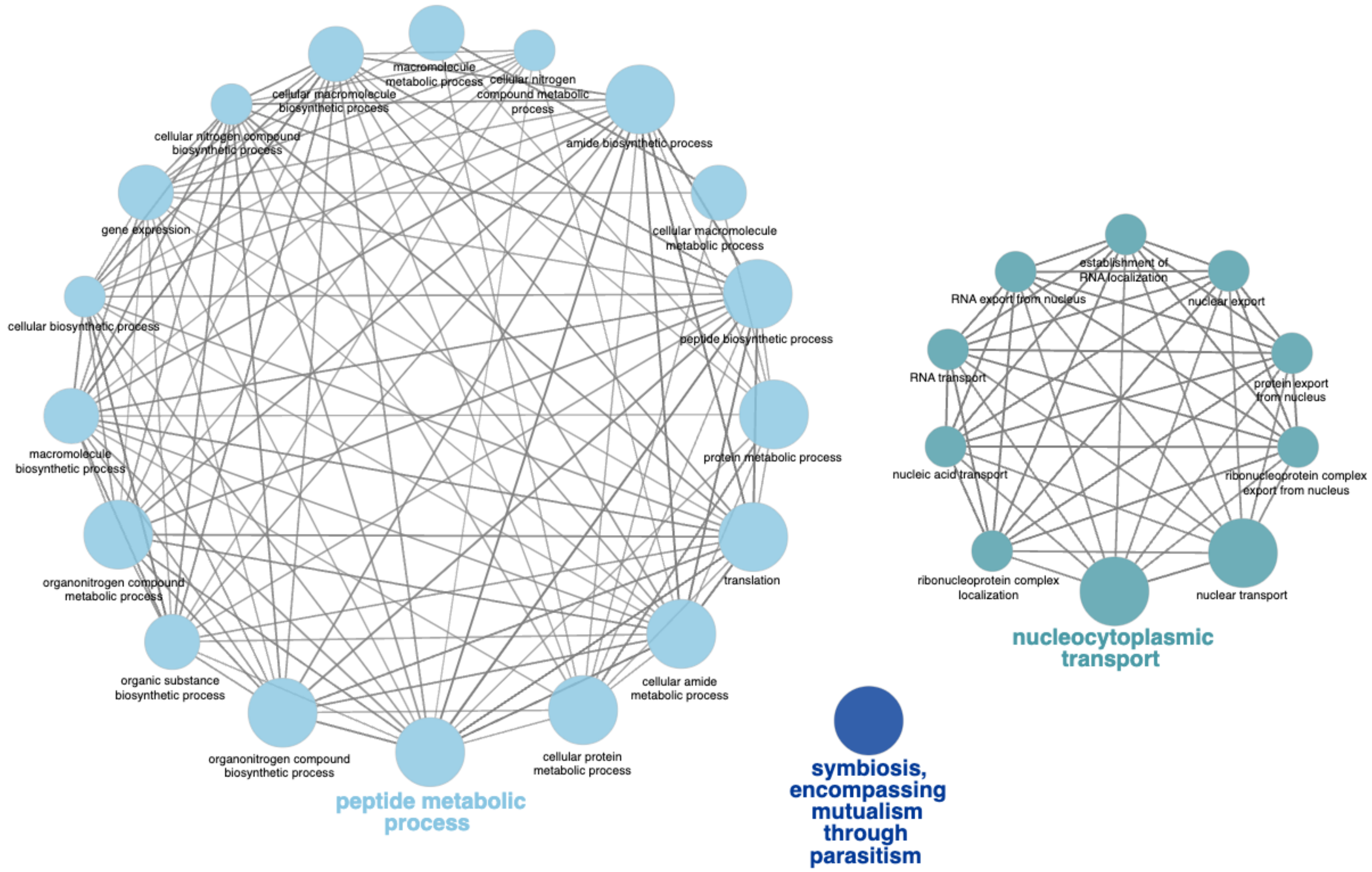


Supplementary Figure-2

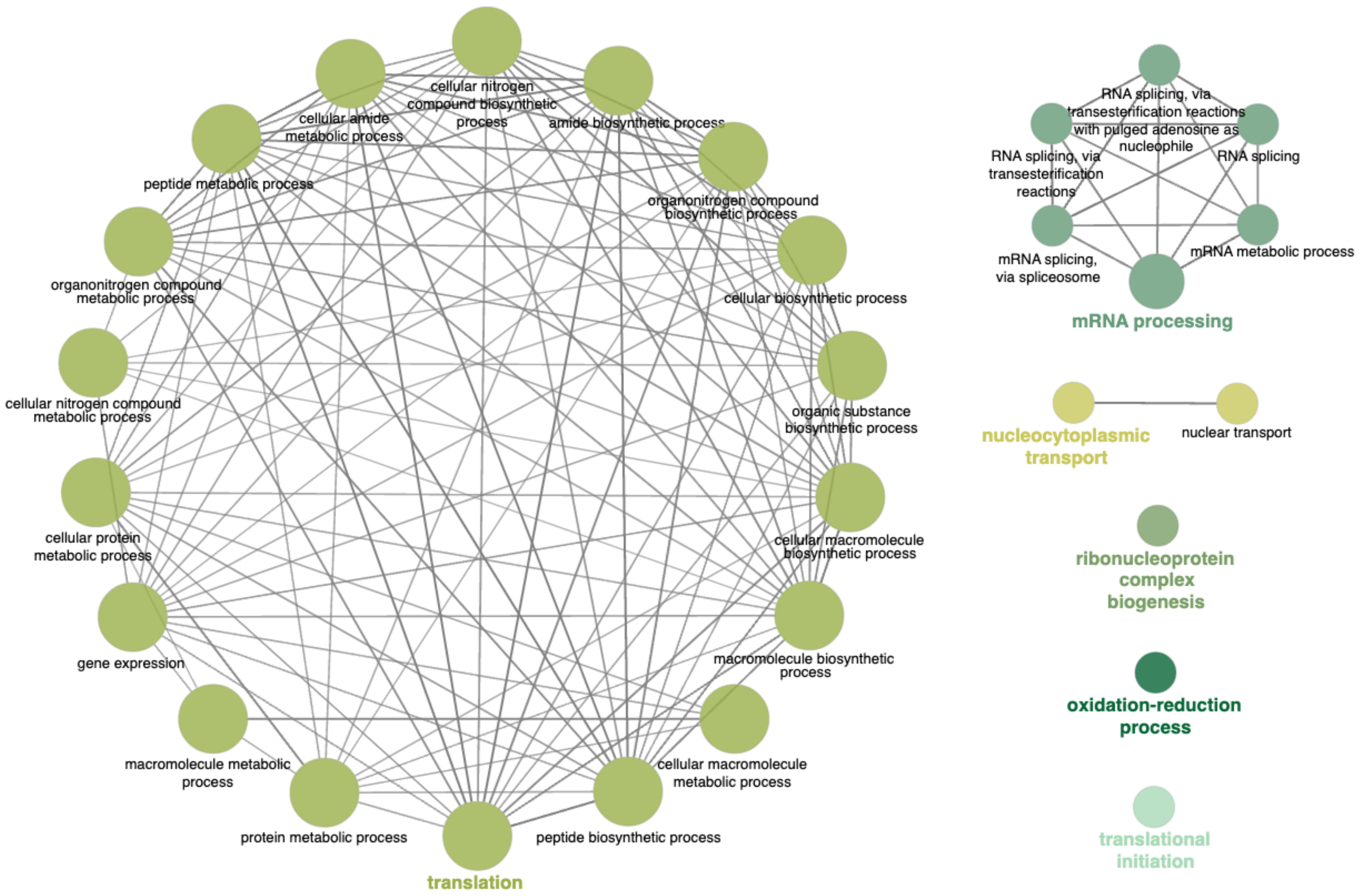


Supplementary Figure-3

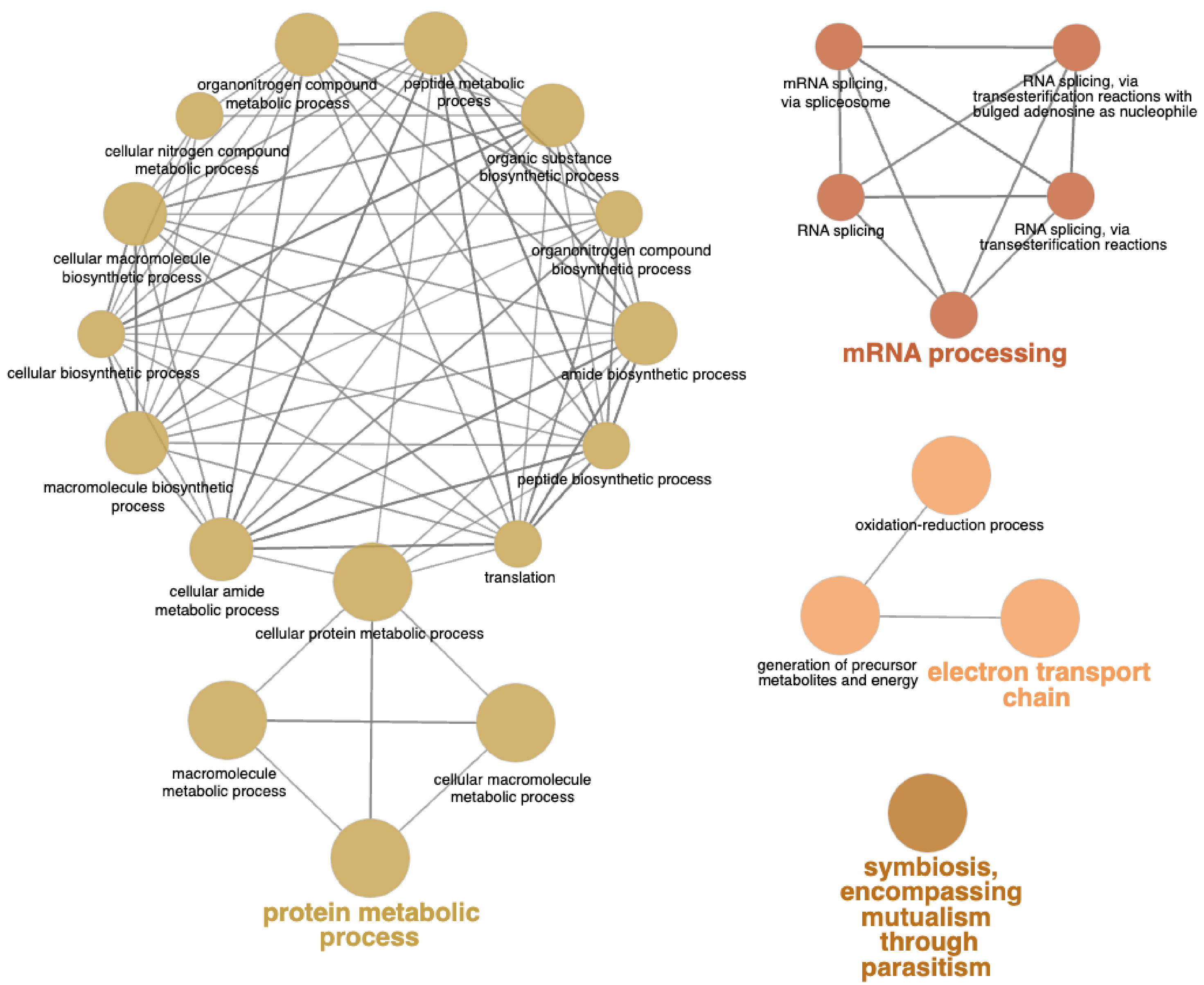


Supplementary Figure-4

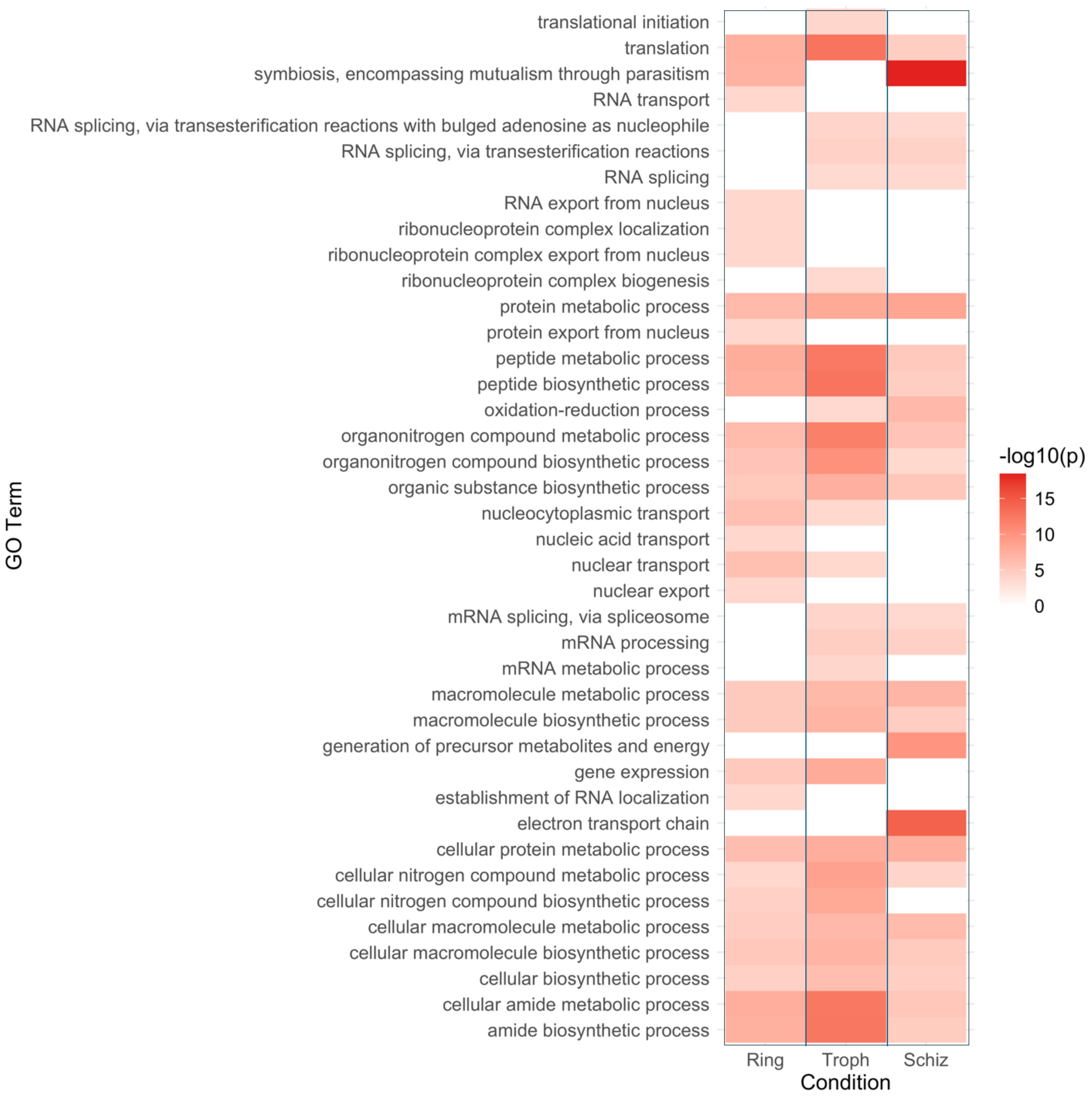
translational initiation
translation
symbiosis, encompassing mutualism through parasitism
RNA transport
RNA splicing, via transesterification reactions with bulged adenosine as nucleophile
RNA splicing, via transesterification reactions
RNA splicing
RNA export from nucleus
ribonucleoprotein complex localization
ribonucleoprotein complex export from nucleus
ribonucleoprotein complex biogenesis
protein metabolic process
protein export from nucleus
peptide metabolic process
peptide biosynthetic process
oxidation-reduction process
organonitrogen compound metabolic process
organonitrogen compound biosynthetic process
organic substance biosynthetic process
nucleocytoplasmic transport
nucleic acid transport
nuclear transport
nuclear export
mRNA splicing, via spliceosome
mRNA processing
mRNA metabolic process
macromolecule metabolic process
macromolecule biosynthetic process
generation of precursor metabolites and energy
gene expression
establishment of RNA localization
electron transport chain
cellular protein metabolic process
cellular nitrogen compound metabolic process
cellular nitrogen compound biosynthetic process
cellular macromolecule metabolic process
cellular macromolecule biosynthetic process
cellular biosynthetic process
cellular amide metabolic process
amide biosynthetic process
GO Term
Ring
Troph
Schiz
Condition
-log10(p)
15
10
5
0

### TEseq pipeline

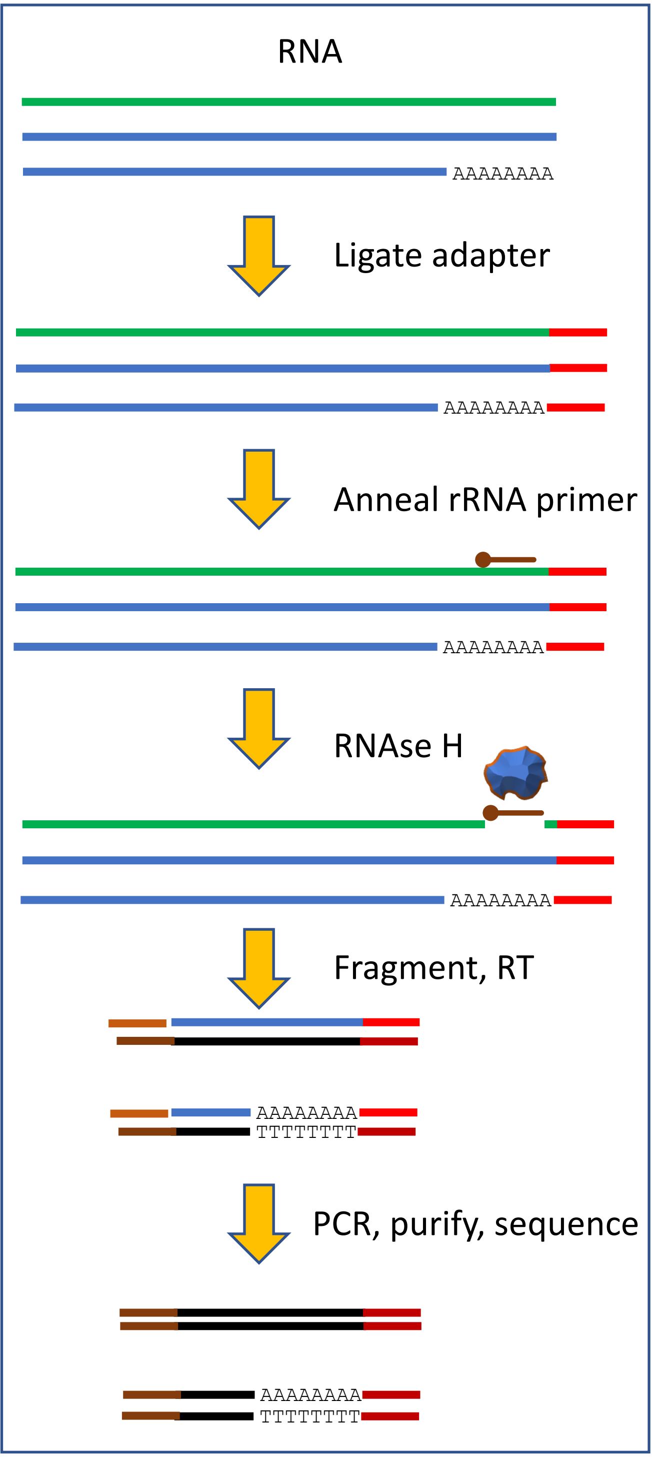


**Supplementary Figure -1**
